# Trophic cooperation promotes bacterial survival of *Staphylococcus aureus* and *Pseudomonas aeruginosa*

**DOI:** 10.1101/2020.06.17.156968

**Authors:** Laura Camus, Paul Briaud, Sylvère Bastien, Sylvie Elsen, Anne Doléans-Jordheim, François Vandenesch, Karen Moreau

**Author notes:** Address correspondence to Karen Moreau, – Université Claude Bernard Lyon1 – CIRI – Pathogénie des staphylocoques – 7 rue Guillaume Paradin – 69 008 Lyon - France –. All authors declare no competing interests.

## Abstract

In the context of infection, *Pseudomonas aeruginosa* and *Staphylococcus aureus* are frequently co-isolated, particularly in cystic fibrosis (CF) patients. Within lungs, the two pathogens exhibit a range of competitive and coexisting interactions. In the present study, we explored the impact of *S. aureus* on the physiology of *P. aeruginosa* in the context of coexistence. Transcriptomic analyses showed that *S. aureus* significantly and specifically affects the expression of numerous genes involved in *P. aeruginosa* carbon and amino acid metabolism. In particular, 65% of the strains presented considerable overexpression of the genes involved in the acetoin catabolic (*aco*) pathway. We demonstrated that acetoin is (i) produced by clinical *S. aureus* strains, (ii) detected in sputa from CF patients, and (iii) involved in *P. aeruginosa’s aco* system induction. Furthermore, acetoin is catabolized by *P. aeruginosa*, a metabolic process that improves the survival of both pathogens by providing a new carbon source for *P. aeruginosa* and avoiding the toxic accumulation of acetoin on *S. aureus*. Due to its beneficial effects on both bacteria, acetoin catabolism could testify to the establishment of trophic cooperation between *S. aureus* and *P. aeruginosa* in the CF lung environment, thus promoting their persistence.

## Introduction

Infectious sites constitute rich microbial ecosystems shared by a large diversity of microorganisms, including the native microbiota and pathogens. Lungs of Cystic Fibrosis (CF) patients are a well-known example of this microbial richness as they gather more than 60 genera of bacteria (1, 2). This density of microorganisms promotes their interactions, through which they model their biological activities and their environment (3, 4). However, microbial interactions are dynamic and range from antagonism to cooperation according to species and environmental conditions (5). For instance, the opportunistic pathogen *Pseudomonas aeruginosa* is well-known for its competitiveness in various ecosystems due to many quorum-sensing-mediated factors such as phenazines, rhamnolipids and the type 6 secretion system (6). *P. aeruginosa* thus can alter the growth, biofilm formation and respiration of yeasts, fungi, and Gram-negative and –positive bacteria (6). Among them, *Staphylococcus aureus* is particularly sensitive to *P. aeruginosa* virulence factors and they can directly lyse staphylococci (7). This competitive interaction between *P. aeruginosa* and *S. aureus* is observed for both environmental and clinical strains as both pathogens are frequently co-isolated from wounds and CF lung samples (7, 8). Until recently, the anti-staphylococcal behavior of *P. aeruginosa* was the only expression observed between the two species and was thus extensively described (7). In the context of CF lung infections, this competitive interaction is highlighted by the decreased prevalence of *S. aureus* as *P. aeruginosa* colonizes lungs during adolescence (9). However, Baldan *et al*. first noted that a non-competitive state between *P. aeruginosa* and *S. aureus* could establish during the development of CF chronic infections (10), calling into question the antagonistic model between the two pathogens.

The establishment of this particular non-competitive state between *P. aeruginosa* and *S. aureus* seems to be linked to the adaptation of *P. aeruginosa* to the pulmonary ecosystem. In fact, selective pressures present in the CF lung environment, such as the host immune system and antibiotic treatments, drive *P. aeruginosa* isolates towards a state of low-virulence and high-resistance (11–14). Major virulence factors involved in quorum-sensing and motility are often mutated in *P. aeruginosa* chronic infection isolates, inducing a decrease in the production of anti-staphylococcal factors followed by non-competitive interaction (15). In addition, the rewiring of metabolism networks and the decrease of its catabolic repertoire also accompany *P. aeruginosa’s* adaptation to the CF environment. This trophic specialization commonly leads to the slower growth of chronic isolates and thus less competitive behavior regarding shared resources (16). Several independent studies observed this state of coexistence between *S. aureus* and *P. aeruginosa* isolated from chronic infections (10, 15, 17, 18). Briaud *et al*. recently demonstrated that this interaction pattern appears to be more frequent than expected. Indeed, among the quarter of CF patients co-infected by both pathogens, 65% were infected by a coexisting *S. aureus-P. aeruginosa* pair (18, 19). Recent studies have shown that coexistence between *P. aeruginosa* and *S. aureus* could promote their persistence throughout the establishment of cooperative interaction. In these conditions, coexisting bacteria demonstrated increased tolerance to antibiotics: to tobramycin and tetracycline for *S. aureus* and to gentamicin for *P. aeruginosa*. This appeared to be related to the induction of small colony variants (15, 17, 18, 20). However, the effects of coexistence on general bacterial physiology, and not only virulence-associated traits, have not yet been explored. Therefore, despite its significance in infectious ecosystems, coexistence between *P. aeruginosa* and *S. aureus* remains poorly understood.

Using global and targeted transcriptomic approaches, we evaluated the impact of the presence of *S. aureus* on the gene expression of *P. aeruginosa* using a set of clinical pairs of strains isolated from CF co-infected patients. Coexistence with *S. aureus* induced the overexpression of many genes involved in the utilization of alternative carbon sources in *P. aeruginosa*, such as amino acids and acetoin. Acetoin was shown to be produced by clinical *S. aureus* isolates *in vitro* and in CF sputum, and catabolized by *P. aeruginosa*. The beneficial effects of acetoin catabolism on both bacteria during their interaction highlighted trophic cooperation between *P. aeruginosa* and *S. aureus* in CF lung infections.

## Materials and Methods

### Bacterial strains

The bacterial strains and plasmids used in this study are listed in Tables S1 and S2. CF clinical strains were isolated by the Infectious Agents Institute (IAI) from sputa of patients monitored in the two CF centres of Lyon (Hospices Civils de Lyon (HCL)). *S. aureus* and *P. aeruginosa* strains were isolated from patients co-infected by both bacteria. Each strain pair indicated in Table S1 was recovered from a single sample, obtained from different patients in most cases, as indicated in Table S1. All the methods were carried out in accordance with relevant French guidelines and regulations. This study was submitted to the Ethics Committee of the HCL and registered under CNIL No 17-216. All the patients were informed of the study and consented to the use of their data.

As schematized in Figure S1A, interaction state was determined for each pair by growth inhibition tests on Tryptic Soy Agar (TSA) and in liquid cultures (18). As previously described, coexistence was characterized by: (i) the absence of inhibition halo on agar tests, and (ii) similar growth in mono and co-cultures for 8 hours (18).

The knock-out Δ*acoR* and Δ*aco* mutants were generated in the *P. aeruginosa* PA2600 strain via allelic exchange thanks to suicide plasmids pEXG2Δ*acoR* and pEXG2Δ*aco* constructed as described previously (21, 22). Gentamicin (Euromedex) was used at final concentrations of 50μg/ml. Detailed protocols are given in supplementary data.

### Cultures conditions

Strains were grown in monoculture or co-culture in Brain-Heart Infusion (BHI), as described by Briaud *et al*. (18). Briefly, *S. aureus, P. aeruginosa, Burkholderia cenocepacia, Stenotrophomonas maltophilia* and *Bacillus subtilis* overnight precultures were diluted to an OD_600_ of 0.1 in BHI and grown for 2h30 at 37°C and 200rpm. Cultures were then diluted to an OD_600_ of 2 in fresh medium and 10ml was mixed with 10ml BHI for monocultures. Co-cultures were performed by mixing 10ml of *P. aeruginosa* suspension with 10ml of *S. aureus*, *B. cenocepacia*, *S. maltophilia* or *B. subtilis* suspension. For supernatant exposure, 10ml of *S. aureus* supernatant from a 4-hour culture was filtered on a 0.22μM filter and added to 10ml of *P. aeruginosa* suspension. Cultures were grown for 8 hours at 200rpm and at 37°C for transcriptomic studies. Long-term survival assays were performed by extending the incubation time of *S. aureus* and *P. aeruginosa* mono- and co-cultures up to five days. Plating at day 0, 3 and 5 was performed on mannitol salt agar (MSA, BBL™ Difco) and cetrimide (Difco™) for *S. aureus* or *P. aeruginosa* counts, respectively. For growth monitoring in the presence of acetoin, minimal medium M63 (76mM (NH_4_)_2_SO_4_, 500mM KH_2_PO_4_, 9μM Fe.SO_4_.7H_2_O, 1mM MgSO_4_.7H_2_O) was inoculated with *P. aeruginosa* to an OD_600_ of 0.1 and incubated for 25h at 37°C and 200rpm. Every two hours during 10 hours and at t=24h, acetoin was added to obtain a final concentration of 1.5mM in the culture. Plating on TSA (Tryptic Soy Agar) was performed at t=0, one hour after each acetoin addition and at t=24h.

### Genome sequencing and annotation

Genome sequencing and annotation of PA2596 and PA2600 strains were performed as previously described (18). In order to compare CDS from PA2596 and PA2600, PAO1 strain (NC_002745.2) was used as a reference. Protein sequences were compared and grouped using a similarity threshold of 95% through Roary (V3.8.2) (23). Gene names and numbers were gathered from PAO1 and used as ID tags for common genes. For non-common genes, CDS from PA2596 and PA2600 were tagged with a specific name (gene number and name recovered from the UniprotKB database). Functional classification was performed using the KEGG database and by a manual literature check of each gene function. The complete genome sequences for the PA2596 and PA2600 strains were deposited in GenBank under the accession number GCA_009650455.1 and GCA_009650545.1.

### Transcriptomic analysis

Figure S1 schematizes the global methodology used for the transcriptomic analysis. RNAseq analysis was performed on four pairs: the patient-specific pairs SA2597/PA2596 (competitive) and SA2599/PA2600 (coexisting), and the crossed pairs SA2599/PA2596 (competitive) and SA2597/PA2600 (coexisting). RNA extraction, cDNA library preparation and sequencing and read treatment were conducted as previously described (18). Gene expression was considered as dysregulated when: (i) the fold change between co-culture and monoculture was at least 4-fold with an adjusted *P*-value<0.05, (ii) dysregulation was observed in the two pairs of strains with the same interaction state, (iii) the dysregulation was specific to either coexistence or competition state. RNAseq data that support our findings are available in the SRAdatabase under the BioprojectID: PRJNA562449, PRJNA562453, PRJNA554085, PRJNA552786, PRJNA554237, PRJNA554233.

Confirmation of gene expression was achieved by RT-qPCR as previously described on 14 or 21 coexisting pairs including the SA2599/PA2600 couple used in the RNAseq experiment **(Fig. S1)** (18). Housekeeping genes *gyrB* and *rpoD* were used as endogenous control. Table S3 lists the primers used and the target genes.

### Acetoin and glucose dosages

Acetoin was quantified using a modified Voges-Proskauer test (24) optimized in 96-well microplates. 35μL of creatine (0.5% m/v, Sigma), 50μL of α-naphtol (5% m/v, Sigma) and 50μL of KOH (40% m/v, Sigma) were added sequentially to 50μL of culture supernatant. The mix was incubated for 15 minutes at room temperature and optical density at 560nm was read using spectrophotometry (Tecan Infinite Pro2000, Tecan-Switzerland). Glucose dosages were performed with a glucose (Trinder, GOPOD) assay kit (LIBIOS) in microplates. 185μL of dosage reagent was added to 5μL of culture supernatant and incubated at 37°C for 5 minutes. Optical density at 540nm was measured. Acetoin and glucose standards (Sigma) were performed in water, BHI or M63 according to the experiment.

### Statistical analysis

Statistical analyses were performed using Prism GraphPad 8.0 software (San Diego, CA). Differences in gene expression fold change and bacterial survival were studied using one-way ANOVA with Dunnett’s or Tukey’s post-test comparisons, as specified in the corresponding figures. Median acetoin and glucose concentrations were compared using Mann-Whitney tests or Kruskall-Wallis tests with Dunn’s correction when appropriate. Differences were considered significant when *P*-values were lower than or equal to 0.05.

## Results

### The *P. aeruginosa* transcriptome is affected by the presence of *S. aureus*

We studied the genetic expression of *P. aeruginosa* in the absence or presence of *S. aureus* in two contexts: when *P. aeruginosa* and *S. aureus* were in competition or when they were in coexistence. We thus performed RNAseq analyses using a competitive strain pair (SA2597/PA2596) and a coexisting one (SA2599/PA2600). Each pair was recovered from a single sample of a co-infected CF patient **(Table S1)**. As the nature of interaction was determined solely by *P. aeruginosa* (18), the pairs were crossed to study gene expression in additional competitive (SA2599/PA2596) and coexisting (SA2597/PA2600) pairs. The transcriptomic effect was therefore evaluated during co-cultures of two competitive and two coexisting strain pairs **(Fig. S1)**. *P. aeruginosa* gene expression was considered dysregulated when dysregulation was common to both co-cultures in comparison to monoculture. Each dysregulated gene was then associated with a functional class by carrying out a KEGG analysis **(Fig. 1A, Tables S4 and S5)**.

**Figure 1:**
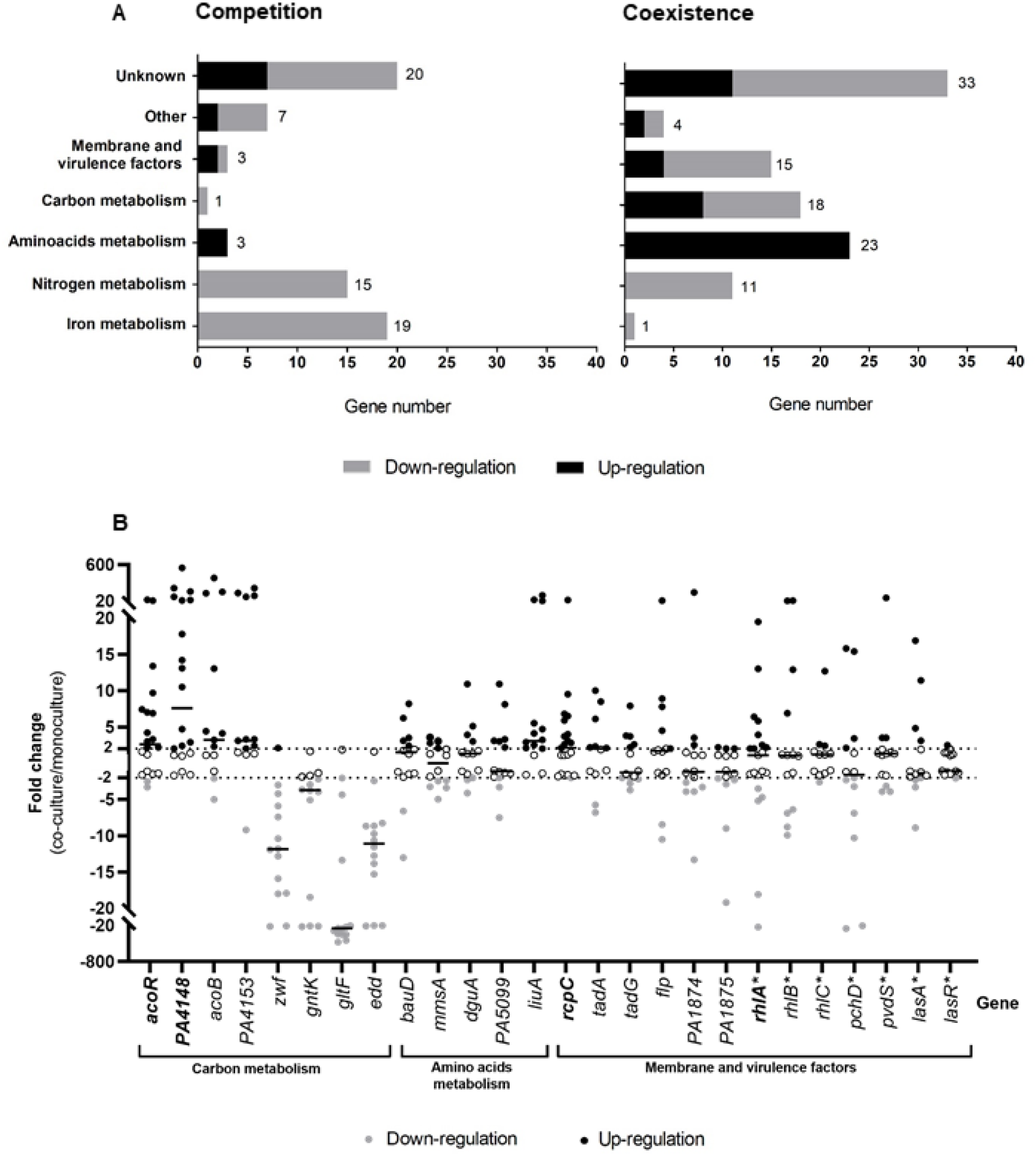
Alteration of the *P. aeruginosa* transcriptome induced by co-culture with *S. aureus*. **A. Number of under-expressed (grey bars) and over-expressed (black bars) genes of *P. aeruginosa* during co-culture with *S. aureus* for competitive (left) or coexisting (right) pairs.** PA2596 competition and PA2600 coexistence strains were cultivated in the absence or presence of SA2597 or SA2599, as described in Fig. S1. RNAs were extracted after 4 hours of culture and the RNAseq analysis was performed as described in the material and methods section. A gene was considered as differentially expressed when the Fold Change (FC) was > |2log_2_| with an adjusted *P*-value <0.05. Functional classification was performed using the KEGG database and the literature. **B. Fold change of 26 *P. aeruginosa* gene expression induced by co-culture with *S. aureus* during coexistence interaction.** Twenty-one coexisting *S. aureus-P. aeruginosa* pairs were isolated from separate CF sputa recovered from 20 different patients. Each *P. aeruginosa* strain was cultivated in the absence or presence of its co-isolated *S. aureus* strain. RNAs were extracted after 4 hours of culture and gene expression was assayed by RT-qPCR. A gene was considered as differentially expressed when the Fold Change (FC) was > |2|, indicated by dotted lines. Black lines indicate the median. Genes were tested in 14 (regular character) or 21 (bold character) strains. The list of the strains used is shown in Table S1. Genes annotated with (*) were not identified as dysregulated during the RNAseq experiment.

Sixty-eight *P. aeruginosa* genes were dysregulated in co-culture in the context of competition, with a majority of down-regulated genes (79.4%, **Fig. 1A**). Fifteen genes involved in nitrogen metabolism and 19 genes involved in iron metabolism were down-regulated, making these two functional classes the most affected in competition. Among these genes, the *nir* and *nos* systems involved in denitrification as well as genes implicated in iron uptake and transport *(isc* and *fec* genes) were down-regulated **(Table S4)**. Other functional classes were less affected, as only 4 genes linked to carbon and amino acid metabolism (*bauA*, *ddaR*, *gntK*, *arcD*) and 3 genes encoding membrane and virulence factors (*rfaD*, *PA2412, cdrA)* were dysregulated in the presence of *S. aureus*.

More genes were affected during coexistence interaction, as the dysregulation of 105 genes was observed in *P. aeruginosa* **(Fig. 1A)**. In spite of a trend of up-regulation (56.4% of genes), we could observe the down-regulation of 11 genes involved in nitrogen metabolism. Among these, 8 were also down-regulated in the context of competition, especially genes from the *nir* system. We can thus presume that the down-regulation of *nir* genes is not specific to coexistence or competition interaction states. Conversely, the dysregulations of iron metabolism related genes appeared to be specific to competition strains, as only one gene (*fhp*) of this functional class was down-regulated in coexistence. However, coexistence specifically affected numerous genes belonging to functional classes of carbon and amino acid metabolism (18 and 23 genes) and membrane and virulence factors (15 genes). Concerning the latter class, a trend towards lower expression was observed, probably due to the down-regulation of membrane associated factors such as the *flp-tad* system *(flp, tad* and *rcpC* genes encoding Flp pilus) and the *PA1874-1876* operon (encoding an efflux pump) **(Table S5)**.

The classes most affected in coexistence with *S. aureus* were related to *P. aeruginosa* energetic metabolism **(Fig. S2)**. We observed a down-regulation of genes coding for major pentose phosphate pathway enzymes, like the gluconokinase GntK, its regulator GntR and the operon *zwf-edaA* (*PA3183-PA3181*), encoding a glucose-6-phosphate 1-dehydrogenase, a 6-phosphogluconolactonase and a 2-keto-3-deoxy-6-phosphogluconate aldolase (25). We also noticed the down-regulation of 5 other genes clustered near this operon and involved in the same pathway (*edd* and *gapA* genes) and glucose transport (*gltB*, *gltF* and *gltK*) (25, 26).

In contrast, an up-regulation of numerous genes involved in the utilization of alternative carbon sources as butanoate and amino acids was observed. The *aco* system, comprising the operon *PA4148-PA4153* and the gene encoding its transcriptional regulator *acoR (PA4147)*, was up-regulated in *P. aeruginosa* coexisting with *S. aureus*. This system has been described in *P. aeruginosa* PAO1 to be responsible for 2,3-butanediol and acetoin catabolism (27). According to KEGG analysis, the up-regulated genes *acsA (PA0887), PA2555 (acs* family) and *hdhA (PA4022)* are also involved in the butanoate pathway and energy production from 2,3-butanediol and acetoin, as their products catalyse the production of acetyl-coA from acetaldehyde and acetate **(Fig. S2)**. Finally, 23 genes implicated in amino acid metabolism were up-regulated in *P. aeruginosa* in the presence of *S. aureus*. Most of them were linked to the catabolism of several amino acids **(Fig. S2)**. In particular, we observed the *liu* operon (*PA2015-PA2012*), the *mmsAB* operon (*PA3569-PA3570*) and the *hut* gene system (*PA5097-PA5100*) involved in leucine, valine and histidine catabolism, respectively.

In order to confirm these transcriptomic effects, we co-cultivated a set of *P. aeruginosa-S. aureus* coexistence CF pairs and performed RT-qPCR to evaluate *P. aeruginosa* gene expression in the presence of *S. aureus* **(Fig. 1B)**. Each pair was isolated from a single sputum. Twenty-six genes were tested, including 19 identified as dysregulated during RNAseq analysis and belonging to the most impacted categories, *i.e*. carbon and amino acid metabolism, and membrane and virulence factors. Most of the genes were tested in a total of 14 *P. aeruginosa-S. aureus* pairs; the expression of four of these genes was assessed in seven additional pairs to confirm the dysregulations observed. The different *P. aeruginosa* strains presented very different transcriptomic patterns during co-cultivation with *S. aureus*, especially for membrane-associated and virulence factor genes. We noticed an over-expression of *rcpC, tadA, tadG* and *flp* from the *flp-tad* system from 52.4% to 35.7% of the *P. aeruginosa* strains. We also tested 7 additional genes encoding virulence factors previously described as involved in *P. aeruginosa* interaction with *S. aureus* as *las*, *rhl, pch* and *pvd* genes (7). Regarding the latter genes, no clear effect of *S. aureus* co-cultivation was observed.

However, clearer transcriptomic patterns were observed for genes linked to carbon and amino acid metabolism, the two gene classes most impacted during the RNAseq experiment **(Fig. 1B)**. We confirmed the up-regulation of *liuA* gene in 78.6% (11/14) of *P. aeruginosa* strains co-cultivated with *S. aureus*. We also confirmed the down-regulation of glucose metabolism genes in a high proportion of strains, ranging from 92.9% (13/14) for *zwf, gltF* and *edd* genes to 71.4% (10/14) for *gntK* gene. Finally, the up-regulation of genes involved in butanoate metabolism was confirmed for *acoR* (57.2%, 12/21), *PA4148* (66.7%, 14/21), *acoB* (57.1%, 8/14) and *PA4153* (64.3%, 9/14), suggesting an impact of co-culture on the whole *aco* system in *P. aeruginosa*. In view of these results, we focused our study on four genes: *liuA* and *zwf*, respectively involved in leucine and glucose catabolism, *PA4148*, the first gene of the *aco* operon and *acoR*, both responsible for acetoin catabolism.

### The *P. aeruginosa aco* system is induced by *S. aureus* acetoin

In order to determine if the transcriptomic dysregulations of *acoR, PA4148, liuA* and *zwf* in *P. aeruginosa* PA2600 are specific to interaction with *S. aureus*, we tested the effect of three other bacterial species: *B. cenocepacia* and *S. maltophilia*, as they are sometimes associated with *P. aeruginosa* in CF patients, and *Bacillus subtilis*, as it produces a large amount of acetoin (28). Dosages in *B. cenocepacia* and *S. maltophilia* monocultures confirmed that these bacteria do not produce acetoin **(Fig. S3)** as previously described (29–31). We observed a down-regulation of *P. aeruginosa zwf* gene expression in all co-cultures in comparison to monocultures **(Fig. 2A)**. Regarding *aco* system genes (*acoR* and *PA4148*) and *liuA* gene, only *B. subtilis* induced levels of overexpression similar to *S. aureus* SA2599. We thus hypothesized that acetoin, produced by these two species during our experiment **(Fig. S3)**, may be the inductor signal for these genes during co-culture with *S. aureus*.

**Figure 2:**
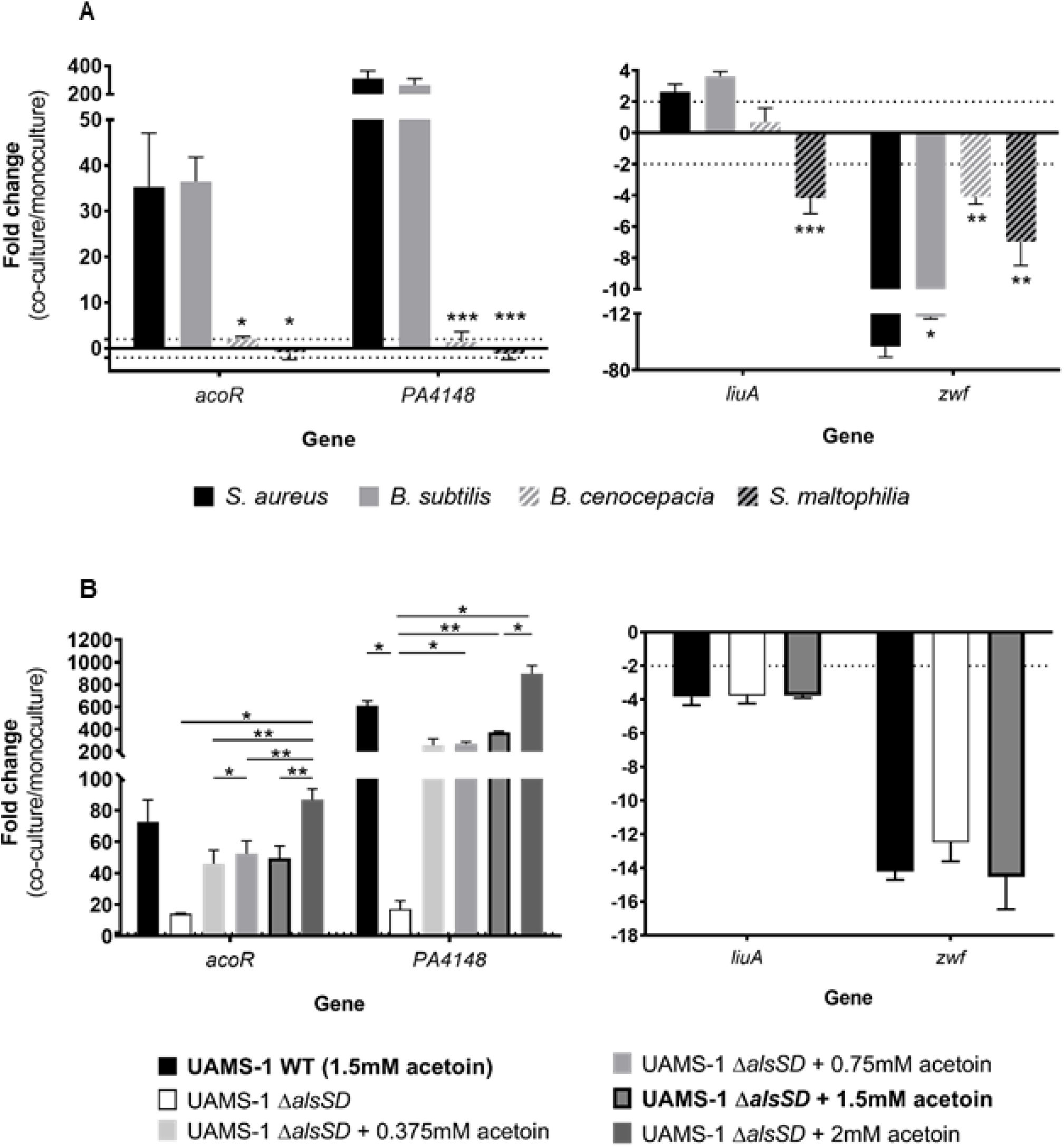
Fold changes of *P. aeruginosa acoR, PA4148, liuA* and *zwf* induced by culture conditions. **A. Fold changes induced by co-culture with *S. aureus* (black bars), *B. subtilis* (grey bars), *B. cenocepacia* (hatched white bars) or *S. maltophilia* (hatched black bars).** *P. aeruginosa* PA2600 strain was cultivated in the absence or presence of *S. aureus* SA2599, *B. subtilis, B. cenocepacia* or *S. maltophilia*. RNAs were extracted after 4 hours of culture and gene expression was assayed by RT-qPCR. Bars represent the mean fold change + SEM from three independent experiments. Dotted lines indicate a fold change = |2|. \**P*_adj_<0.05, \*\**P*_adj_<0.01, \*\*\**P*_adj_<0.001 ANOVA with Dunnett’s correction (*S. aureus vs*. condition). **B. Fold changes induced by culture in supernatant of *S. aureus* UAMS1 wild type (WT, black bars), Δ*alsSD* mutant (white bars), Δ*alsSD* mutant complemented with increasing acetoin concentrations (grey bars).** *P. aeruginosa* PA2600 strain was cultivated in the absence or presence of filtered supernatant from *S. aureus* UAMS-1 WT, Δ*alsSD* or Δ*alsSD* complemented with acetoin concentrations ranging from 0.375mM to 2mM. RNAs were extracted after 4 hours of culture and gene expression was assayed by RT-qPCR. Bars represent the mean fold change + SEM from three independent experiments. The dotted line indicates a fold change = |2|. \**P*_adj_<0.05, \*\**P*_adj_<0.01 ANOVA with Tukey’s correction.

In order to test this hypothesis, we first explored whether an inductor signal was present in the supernatant of *S. aureus* culture. We indeed observed an overexpression of *acoR* and *PA4148* when *P. aeruginosa* PA2600 was cultivated in *S. aureus* SA2599 culture supernatant, as well as the down-expression of *zwf* but to a lesser extent in comparison to co-culture **(Fig. S4)**. On the contrary, we did not observe overexpression of the *liuA* gene, suggesting that this effect is not due to acetoin and requires the presence of *S. aureus* cells. Secondly, we cultivated *P. aeruginosa* PA2600 in the presence of *S. aureus* UAMS-1 WT supernatant or its *ΔalsSD* derivative defective in acetoin synthesis (32). The supernatant of the wild-type UAMS-1 strain induced the same transcriptomic patterns on *P. aeruginosa* as those obtained with the CF SA2599 strain **(Fig. S4)**. In the presence of UAMS-1 *ΔalsSD* supernatant, the overexpression of *aco* genes was almost totally eliminated **(Fig. 2B)**. However, the addition of acetoin to this supernatant restored this overexpression in a dose-dependent manner but with threshold effects between 0.375mM and 1.5mM of acetoin **(Fig. 2B)**. This indicates that induction through acetoin is one of the mechanisms involved in *aco* system overexpression in *P. aeruginosa*. This experiment showed that *zwf* gene down-regulation did not seem to be mediated by acetoin.

### Coexisting isolates of *S. aureus* and *P. aeruginosa* efficiently metabolize acetoin

As the *aco* system is involved in acetoin catabolism in *P. aeruginosa* (27) and *S. aureus* produces this molecule (27, 28), we hypothesized that *P. aeruginosa* could catabolize *S. aureus* acetoin. To explore this hypothesis, we monitored acetoin concentration during mono- and co-cultures of three pairs of strains: SA2599/PA2600 **(Fig. 3A)**, SA146/PA146 and SA153/PA153A **(Fig. S5A and B)**. *P. aeruginosa* strains did not produce acetoin, but acetoin accumulation up to 1.3mM was observed in *S. aureus* monocultures. Interestingly, a reduction of acetoin accumulation of at least 30% was observed when *S. aureus* was co-cultivated with *P. aeruginosa*, in comparison to *S. aureus* monoculture. The same trend could be observed in CF patient sputa, as a higher acetoin concentration was detected in sputa from *S. aureus* mono-infected patients (n=9) in comparison to sputa from *S. aureus*-*P. aeruginosa* co-infected patients (n=11) **(Fig. S6)**. This effect could be due to a down-regulation of *S. aureus* acetoin biosynthesis, or acetoin catabolism in co-culture. Growth of *P. aeruginosa* PA2600 in *S. aureus* SA2599 supernatant containing acetoin actually led to a reduction of acetoin concentration, demonstrating the ability of *P. aeruginosa* to catabolize acetoin **(Fig. 3B)**. This ability was confirmed for the two other strains PA146 and PA153A when grown in the supernatant of SA146 and SA153, respectively **(Fig. S5C and D)**. We also noted that acetoin catabolism started after glucose depletion in the supernatant and could be delayed by glucose addition **(Fig. 3B)**. This suggests that *P. aeruginosa* uses acetoin as an alternative carbon source in the absence of easily available substrates such as glucose. The early glucose depletion observed during co-culture supports this hypothesis **(Fig. 3A)**.

**Figure 3:**
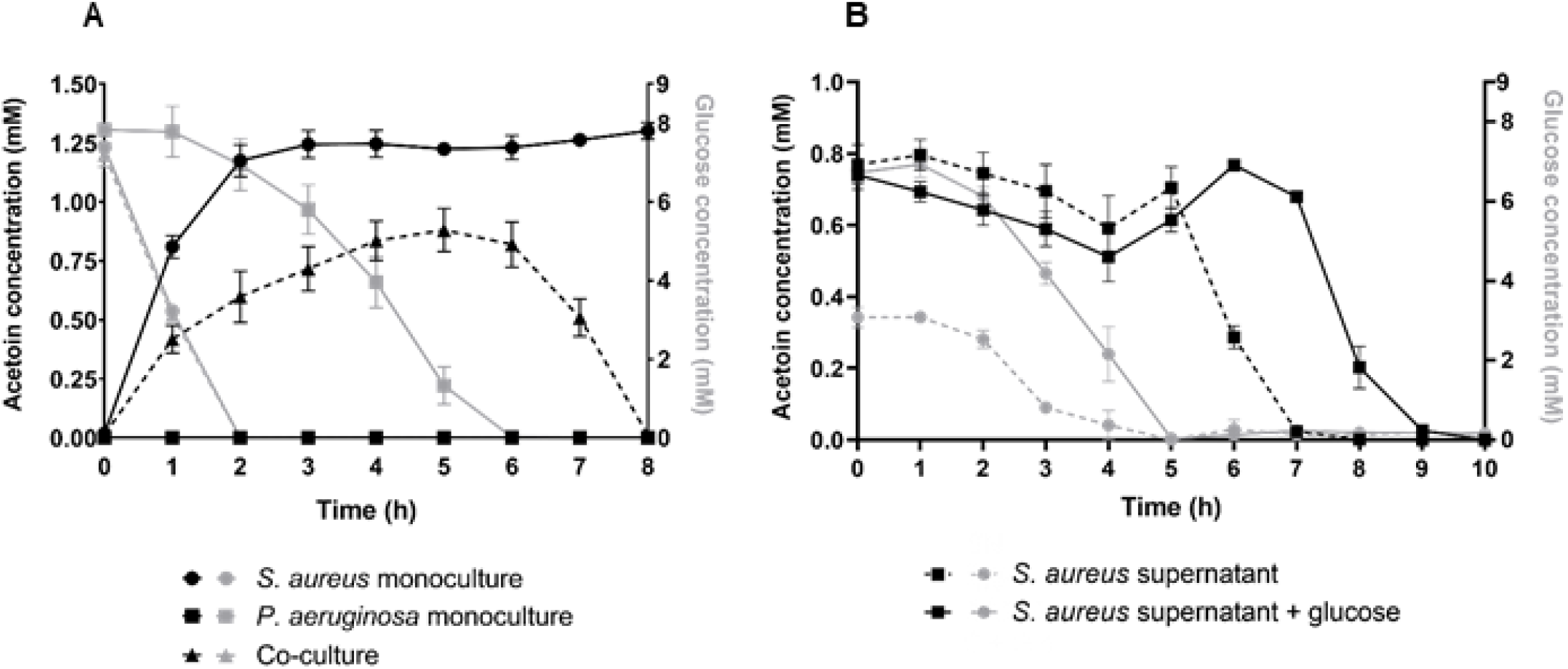
Monitoring of acetoin (black lines) and glucose (grey lines) concentrations in *S. aureus* and *P. aeruginosa* monocultures or co-culture (A) or in *S. aureus* supernatant inoculated with *P. aeruginosa* (B), during coexisting interaction. **A.** *S. aureus* SA2599 and *P. aeruginosa* PA2600 were cultivated in monoculture or co-culture. Acetoin and glucose were quantified from the supernatant each hour. Points represent the mean acetoin or glucose concentration ± SEM from three independent experiments. Similar acetoin accumulation results are shown in Figure S5 for couples 146 and 153A. **B.** A supernatant of *S. aureus* SA2599 filtered for hours was inoculated with *P. aeruginosa* PA2600 culture or sterile medium for controls. The supernatant was used unaltered (dotted lines) or complemented with glucose (solid lines). Acetoin and glucose were quantified from the supernatant each hour. Points represent the mean acetoin or glucose concentration ± SEM from three independent experiments. Similar results of acetoin catabolism are shown in Figure S5 for couples 146 and 153A.

In order to test if this acetoin metabolism was specific to the interaction state between *S. aureus* and *P. aeruginosa*, we evaluated acetoin production and catabolism for 12 couples of competition and 12 couples of coexistence **(Table S1)**. Cultivated in *P. aeruginosa* supernatant, *S. aureus* strains from coexisting pairs were able to produce 4-times more acetoin (230μM) than strains from competitive pairs (60μM) **(Fig. 4A)**. This distinction was not observed during culture in rich medium **(Fig. S7)**. Regarding *P. aeruginosa*, we cultivated the sets of competitive and coexisting *P. aeruginosa* strains in SA2599 supernatant and monitored acetoin catabolism **(Fig. 4B)**. Both competitive and coexisting strains catabolized acetoin since we observed a decrease in acetoin concentration for both groups. However, coexisting strains showed increased catabolism efficiency. Indeed, this group catabolized 98.6% of acetoin after 4 hours of culture, while competitive strains catabolized only 47% of acetoin. This increased the efficiency of acetoin production although catabolism for coexisting strains could not be explained by a difference in glucose utilization between competition and coexistence strains, as both groups catabolized glucose with the same efficiency **(Fig. S8)**. Acetoin production by *S. aureus* and catabolism by *P. aeruginosa* therefore seems to be more efficient in isolates from coexisting couples.

**Figure 4:**
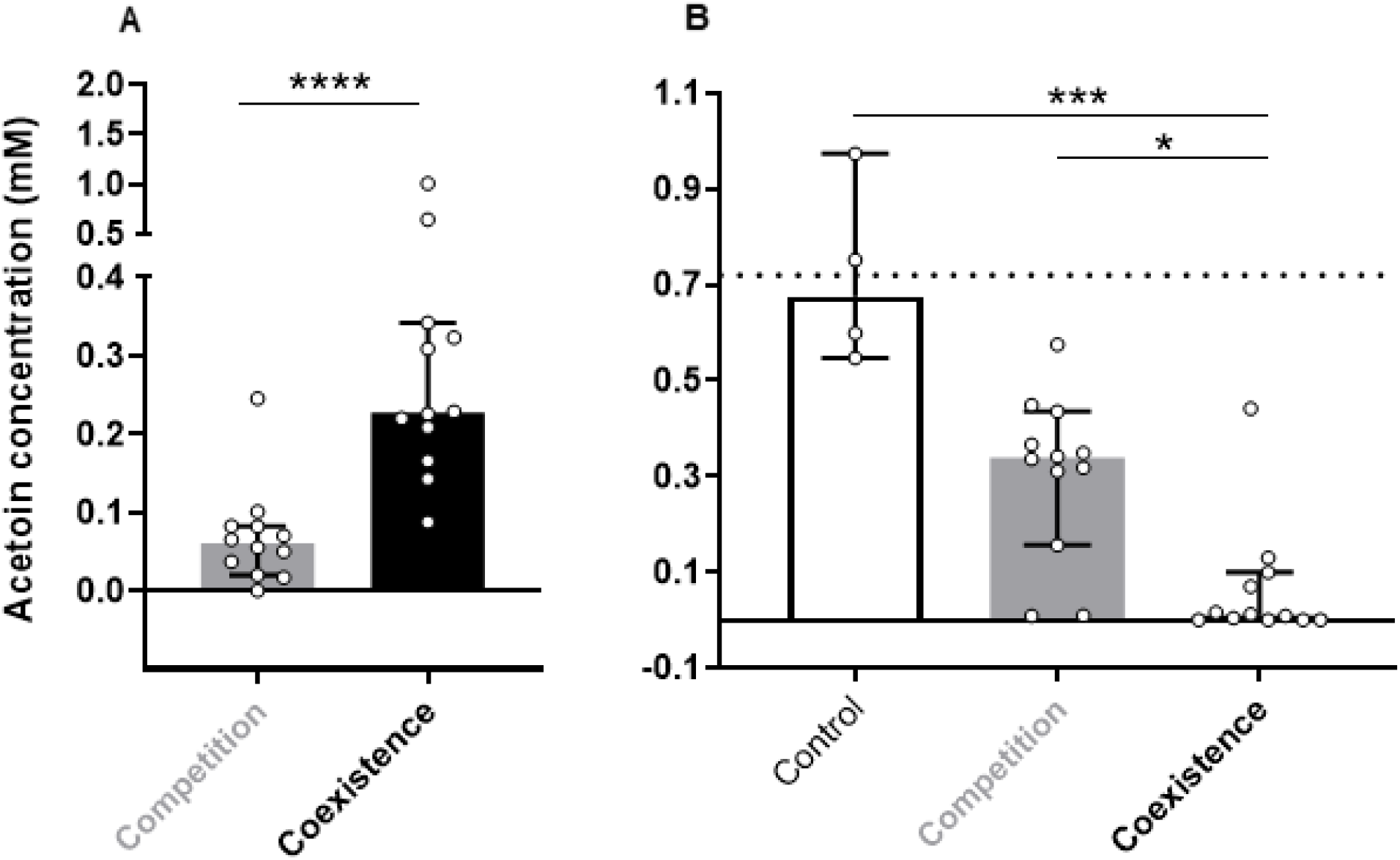
Ability of acetoin production by *S. aureus* (A) and catabolism by *P. aeruginosa* (B) for competitive (grey bars) and coexisting (black bars) strains. **A. Acetoin concentration in *P. aeruginosa* supernatant inoculated with *S. aureus* strains from competition and coexistence couples.** Each *S. aureus* strain from competition (n=12) and coexistence (n=12) couples was cultivated in *P. aeruginosa* PA2600 filtered supernatant and acetoin was quantified from the supernatant after 6 hours of culture. Bars represent the median acetoin concentration ± 95% CI. \*\*\*\**P*<0.0001 Mann-Whitney test. **B. Acetoin concentration in *S. aureus* supernatant inoculated with *P. aeruginosa* strains from competition and coexistence couples.** Each *P. aeruginosa* strain from competition (n=12) and coexistence (n=12) couples was cultivated in *S. aureus* SA2599 filtered supernatant and acetoin was quantified from the supernatant after 4 hours of culture. Sterile supernatant was used as control for acetoin degradation. Bars represent the median acetoin concentration ± 95% CI. The dotted line indicates the initial acetoin concentration. \**P*_adj_<0.05, \*\**P*_adj_<0.01 Kruskall-Wallis test with Dunn’s correction.

### Acetoin catabolism by *P. aeruginosa* increases survival rates of both pathogens in co-culture

As *P. aeruginosa* catabolizes acetoin when medium is glucose-depleted, we tested the effect of acetoin on PA2600 growth in minimal medium M63 (containing no glucose or amino acids) supplemented with 1.5mM acetoin every 2 hours **(Fig. 5)**. *P. aeruginosa* was able to grow up to 1.6×10^9^ CFU/ml after 24h of culture with acetoin as sole carbon source while PA2600 Δ*acoR* and Δ*aco* mutants grew significantly less, reaching a maximum cell concentration of 1.5×10^8^ UFC/ml at 24h **(Fig. 5A)**. In parallel, we quantified acetoin concentration. We observed an accumulation of acetoin throughout the experiment in the presence of PA2600 Δ*acoR* and Δ*aco* mutants. For the WT strain, the accumulation of acetoin reached its maximum at 5h of culture and afterwards slowly decreased to reach undetectable values at 10h of culture, demonstrating the consumption of acetoin in such conditions **(Fig. 5B)**. A delay of 3h between the start of acetoin catabolism and the increased growth of the WT strain was noticed, possibly due to the adaptation of metabolism. Acetoin catabolism nonetheless promoted a 10-fold increase in growth of *P. aeruginosa* during extended culture in glucose depleted medium.

**Figure 5:**
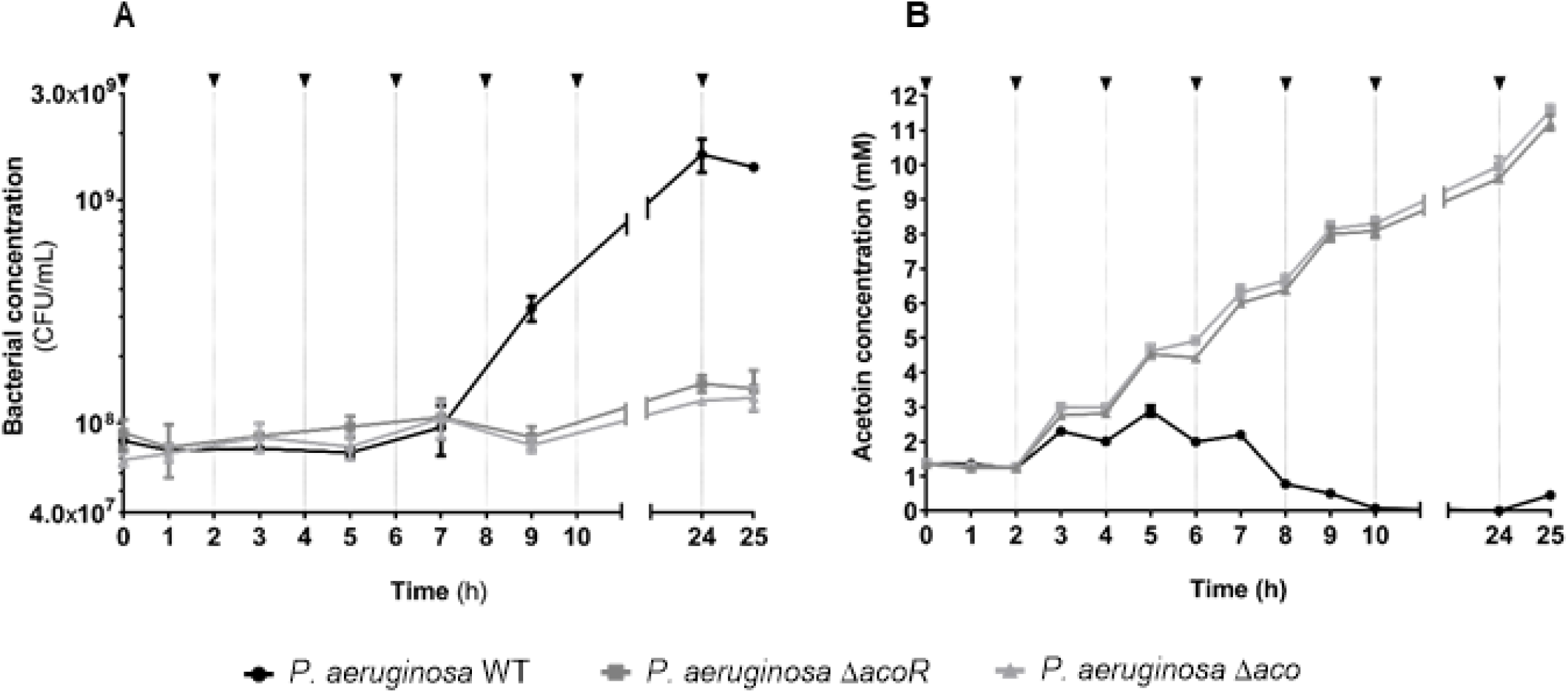
Monitoring of *P. aeruginosa* growth (A) and acetoin concentration (B) in minimal medium supplemented with acetoin. *P. aeruginosa* PA2600 WT, Δ*acoR* and Δ*aco* strains were grown in M63 medium and 1.5mM acetoin was added every 2 hours, indicated by black arrows. **A.** Cultures were plated on TSA each 2 hours to count bacteria. Points represent the mean bacterial concentration ± SEM from three independent experiments. **B.** Acetoin was quantified from the supernatant each hour. Points represent the mean acetoin concentration ± SEM from three independent experiments.

In order to assess the impact of acetoin catabolism on the survival of both pathogens, we co-cultivated *P. aeruginosa* PA2600, *ΔacoR* and *Δaco* mutants with *S. aureus* SA2599. As *S. aureus* was not able to grow in M63 poor medium, and since acetoin affects its long term survival (33, 34), we performed a long term culture (5 days) in BHI medium. We determined the survival rate of *S. aureus* co-cultivated with *P. aeruginosa* in comparison to monoculture **(Fig. 6A)**. Co-culture with the WT strain of *P. aeruginosa* induced a *S. aureus* survival rate of 4.7×10^-1^ after 3 days of culture and of 5.1×10^-3^ after 5 days*. S. aureus* survival thus appears to be highly affected by long-term co-culture with *P. aeruginosa*, even if the strains coexist stably during shorter spans of culture (18). The state of interaction with *S. aureus* thus seems to rely on nutritional conditions, a fact already observed elsewhere (35, 36). However, the survival rate of *S. aureus* was even lower during co-culture with *P. aeruginosa* Δ*acoR* and Δ*aco* mutants, reaching only 9.7×10^-2^ at 3 days of culture and 5.6×10^-4^ at 5 days. *S. aureus* survival was thus 4 to 10 times lower when *P. aeruginosa* was not able to catabolize acetoin, in comparison to co-culture with a strain that catabolize the molecule efficiently. These results suggest that the accumulation of acetoin may impact *S. aureus* survival during co-culture with *P. aeruginosa*. In parallel, we determined the survival rate of *P. aeruginosa* **(Fig. 6B)**. We noticed that in such conditions, the *P. aeruginosa* WT strain presented a growth advantage in co-culture, with a maximum 6.6-fold increase of the *P. aeruginosa* population after 5 days in the presence of *S. aureus* in comparison to monoculture **(Fig. 6B)**. The opposite effect was observed for *aco* mutant strains, as their populations were reduced by 75% for Δ*acoR* and 32% for Δ*aco* after 3 days of co-culture in comparison to monoculture, confirming the role of acetoin catabolism in favouring *P. aeruginosa* growth and survival.

**Figure 6:**
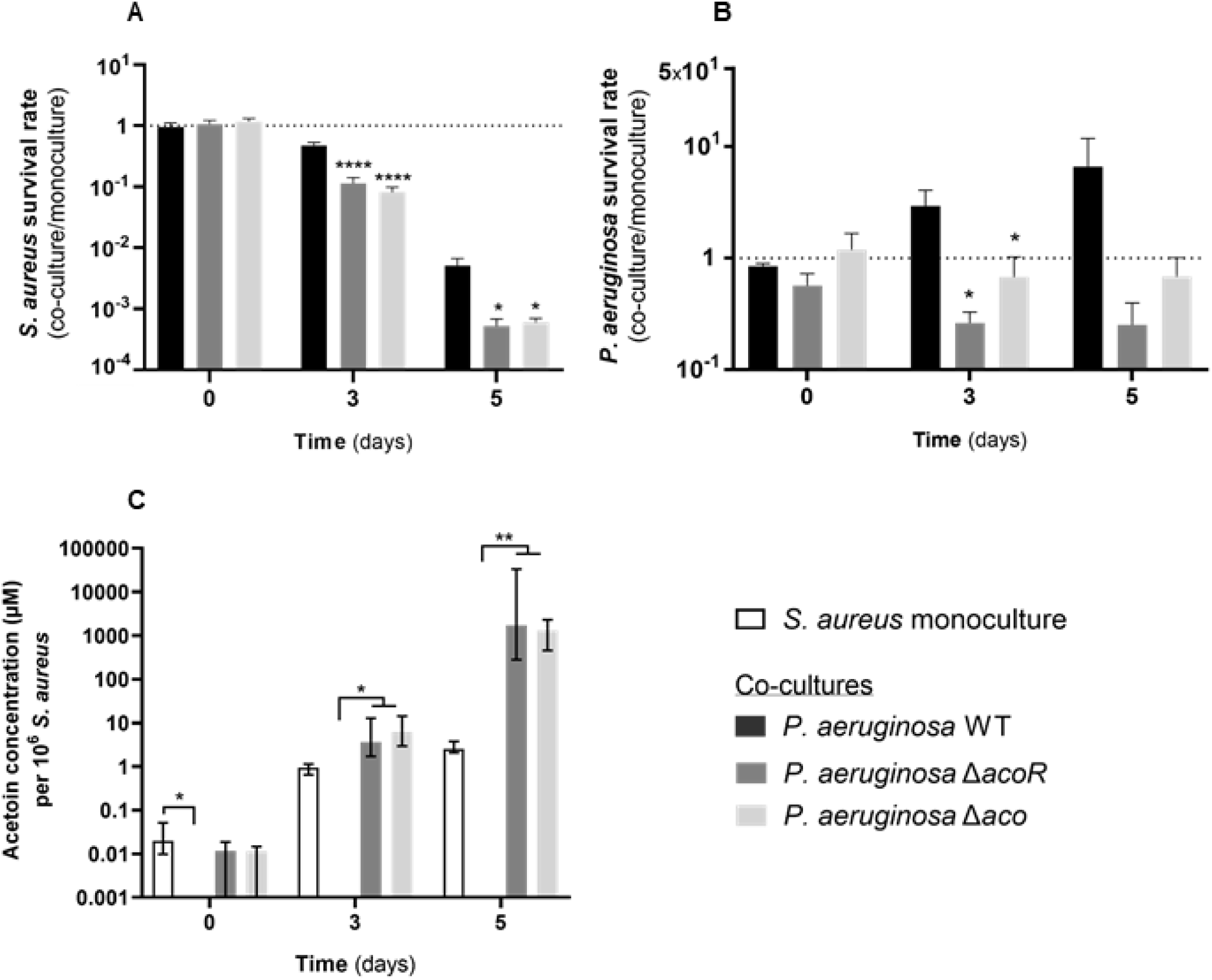
Monitoring of *S. aureus* survival (A), *P. aeruginosa* survival (B) and acetoin concentration (C) during long-term co-culture. *S. aureus* SA2599 was cultivated in the presence of *P. aeruginosa* PA2600 WT, Δ*acoR* and Δ*aco* during 5 days. Cultures were plated at J0, J3 and J5 on MSA and cetrimide to count *S. aureus* and *P. aeruginosa* respectively and acetoin was quantified from supernatant. **A, B.** Survival rate was estimated by dividing the bacterial concentration in co-culture by bacterial concentration in monoculture for each bacterium. Bars represent the mean survival rate *+* SEM from five independent experiments. \**P*_adj_<0.05, \*\*\*\**P*_adj_<0.0001 one-way ANOVA with Dunnett correction (WT *vs*. condition). **C.** Acetoin concentration was normalized to *S. aureus* counts. Bars represent the median acetoin concentration per 10^6^ *S. aureus* ± 95% CI from five independent experiments. \**P*_adj_<0.05, \*\**P*_adj_<0.01, Kruskal-Wallis with Dunn’s correction.

In order to figure out if acetoin accumulation was involved in the decreased survival rate of *S. aureus*, we monitored acetoin concentration in *S. aureus* monoculture and co-cultures **(Fig. 6C)**. As expected, we did not detect acetoin in co-culture with the PA2600 WT strain over the five days but we observed an accumulation of acetoin with PA2600 Δ*acoR* and Δ*aco* mutants. The proportion of acetoin in co-culture (1300μM to 1700μM/10^6^ *S. aureus*) was more than 500 times higher than in monoculture (2.5μM/10^6^ *S. aureus*) **(Fig. 6C)**. We thus cultivated *S. aureus* in different acetoin/cells proportions and observed that acetoin had an inhibitory dose-dependent effect on *S. aureus* growth from 20μM/10^6^ *S. aureus* **(Fig. S9)**. We concluded that acetoin accumulation may be responsible for decreased *S. aureus* survival during co-culture with PA2600 Δ*acoR* and Δ*aco* mutants.

Taken together, these results demonstrated that acetoin catabolism improves *S. aureus* survival in co-culture, in comparison to co-culture with strains unable to catabolize acetoin, since high concentration of acetoin appears to impair *S. aureus* growth. In parallel, acetoin catabolism promotes *P. aeruginosa* survival as a nutritional alternative carbon source.

## Discussion

Coinfection with *S. aureus* and *P. aeruginosa* is a frequent situation, especially in the lungs of CF patients, where coinfection accounts for 25% to 50% of cases (17–19). In this context of coinfection, two states of interaction between the two pathogens have been described: the well-known competitive interaction where *P. aeruginosa* is able to inhibit the growth of *S. aureus* and the coexistence state where the growth of both species is not affected by either individual species. The first state was studied extensively and the leading bacterial determinants of *S. aureus* growth inhibition were described (7). On the contrary, little is known about the impact of the state of coexistence on bacterial physiology. In the present study, we explored the impact of *S. aureus* on *P. aeruginosa* gene expression and physiology.

Comparing competitive and coexistence states, we observed that the down-regulation of the genes involved in iron metabolism was specific to competition. Most of these, such as *fec* genes and *PA4467-PA4471* operon, are involved in ferrous iron uptake and down-regulated during iron-replete conditions (37–39). These conditions are certainly generated by the lysis of *S. aureus* that provides an iron source to *P. aeruginosa* during competitive interaction (7, 36), a situation not observed in coexistence. This hypothesis is supported by the work of Tognon *et al*. (40) that also identified the down-regulation of iron metabolism genes during competitive interaction. Interestingly, they also noted typical responses of amino acid starvation including the down-regulation of genes involved in branched-chain amino acid degradation in competitive *P. aeruginosa* (40). While we did not identify such dysregulation in competition, an overexpression of numerous genes involved in amino acid catabolism was noted during coexistence **(Table S5)**, emphasizing that these dysregulations depend on interaction.

More interestingly, we observed that both carbon and amino acid metabolism was specifically affected during coexisting interaction. Many genes involved in glucose catabolism were down-regulated in coexisting isolates during co-culture with *S. aureus*, especially when the medium was glucose-depleted **(Fig. 3)**. It is noteworthy that the *zwf* gene, down-regulated in almost all the *P. aeruginosa* strains tested, encodes a glucose-6-phosphate dehydrogenase that converts glucose-6-phosphate to 6-phosphogluconate; the first enzyme in the Entner-Doudoroff pathway, which is central to carbon metabolism in *Pseudomonas sp*.

In this condition of glucose depletion, we demonstrated that *P. aeruginosa* was able to use an alternative carbon source provided by *S. aureus:* acetoin. Acetoin is a four-carbon molecule produced by the decarboxylation of α-acetolactate. Owing to its neutral nature, the production and excretion of acetoin during exponential growth prevents over acidification of the cytoplasm and the surrounding medium. When other carbon sources are exhausted, it can constitute an external energy source for fermentive bacteria (28).

Acetoin produced by *S. aureus* was shown to be an inductor of the *aco* operon and *acoR* expression in *P. aeruginosa* **(Fig. 2B)** (27), allowing acetoin catabolism. This occurred in the absence of glucose and was potentially mediated by carbon catabolic repression **(Fig. 3)**, a situation that was already described in other bacteria such as *B. subtilis* (28). However, threshold effects in acetoin-mediated induction and variability in *aco* system overexpression in *P. aeruginosa* strains **(Fig. 1B and 2B)** suggest that other regulatory mechanisms may be involved. Our study may also support the relationship between acetoin and branched-chain amino acid pathways. Indeed, the biosynthesis pathways of acetoin and leucine are co-regulated and share the same precursor α-acetolactate in *S. aureus* (28). In response to co-culture with *S. aureus, P. aeruginosa* clinical strains showed overexpression of acetoin and leucine catabolism genes **(Fig. 1B and 2)**, suggesting the presence of both compounds in our co-culture conditions. All of our analyses were performed *in vitro*. However, using Voges-Proskauer dosage, we were able to confirm the presence of acetoin in CF patient sputa, and in lower concentrations for *P. aeruginosa-positive* samples **(Fig. S6)**. No direct correlation between the presence of *P. aeruginosa* presence and the quantity of acetoin can be established as other microorganisms present in sputa may also have an impact on acetoin concentration. Nevertheless, our data support the work of Španěl *et al*. (41) and suggest that *P. aeruginosa* may catabolize and use *S. aureus* acetoin in the lung environment.

More importantly, we observed that the catabolism of acetoin by *P. aeruginosa* and acetoin production by *S. aureus* were both more efficient for coexisting isolates, in comparison to competitive ones **(Fig. 4)**. This underlines the adapted metabolic regulation in coexisting isolates in comparison to competitive ones. It is well known that the coexistence phenotype between *P. aeruginosa* and *S. aureus* is a consequence of an adaptation process. Indeed, *P. aeruginosa* strains isolated from early infection outcompete *S. aureus* while strains isolated from chronic infection are less antagonistic and can be co-cultivated with *S. aureus* (7, 10, 17). It has also been widely described how both pathogens evolve during colonization to evade the immune response and antibiotic treatment (42). Here, for the first time, we suggest that an evolutionary process leads to an adaptation of interspecies metabolic pathways between *P. aeruginosa* and *S. aureus*.

Therefore, we suggest that acetoin produced by *S. aureus* could contribute to sputum nutritional richness and be used by *P. aeruginosa* to survive in this nutritionally competitive environment during chronic infection. This hypothesis is supported by the beneficial effect of acetoin catabolism on *P. aeruginosa* growth and survival, especially during co-culture with *S. aureus* **(Fig. 5 and 6B)**. *S. aureus* survival in the presence of *P. aeruginosa* was also shown to be highly affected by nutrient availability induced by co-culture conditions **(Fig. 6A)**. While coexistence is characterized by an absence of *S. aureus* growth inhibition during 8-hour co-culture (18), it appears that nutritional competition can still occur under unfavourable conditions and affect *S. aureus’s* survival. Therefore, coexistence between the two pathogens is promoted in nutritionally rich environments, in line with previous observations (35, 36). Although its survival rate is affected under adverse nutritional conditions induced by long-term culture, acetoin catabolism benefits its producer, *S. aureus* **(Fig. 6A)**. Although this effect seems to be linked to acetoin accumulation in the medium, as demonstrated by *P. aeruginosa* Δ*acoR* and Δ*aco* mutants that do not catabolize acetoin anymore **(Fig. 5B and 6C)**, the precise mechanism remains unclear. In *S. aureus*, cell death in the stationary phase may be induced by acetate production and ensuing intracellular acidification. Thomas *et al*. showed that acetoin production counters cytoplasmic acidification by consuming protons and promotes *S. aureus’s* survival in the late-stationary phase (34). We hypothesize that acetoin accumulation in the medium may induce a negative control of acetoin synthesis, affecting *S. aureus’s* survival during co-culture conditions.

Previous studies demonstrated the potential benefits of *S. aureus* and *P. aeruginosa* during coinfection. For example, *S. aureus* facilitates the survival of *P. aeruginosa lasR* mutants commonly found in CF patients by detoxifying surrounding nitric oxide released by host immune cells (43). On the other hand, 4-hydroxy-2-heptylquinoline-N-oxide (HQNO) produced by *P. aeruginosa* cells inhibits respiration in *S. aureus* and also protects *S. aureus* cells from aminoglycosides (44). Additionally, we recently demonstrated that *S. aureus* antibiotic resistance and internalization into epithelial cells were increased in the presence of coexisting *P. aeruginosa* (18). Here, we show that carbon metabolism is largely affected and that *P. aeruginosa* uses the acetoin produced by *S. aureus* as an alternative carbon source. This metabolic dialogue between the two pathogens is selected during bacterial adaptation in CF lungs and promotes their survival. Thus, we highlight for the first time trophic cooperation between *S. aureus* and *P. aeruginosa* during cooperative interaction.

## Acknowledgments

This work was supported by the Fondation pour la Recherche Médicale, grant number ECO20170637499 to LC; the Finovi foundation to KM; the associations “Vaincre la mucoviscidose” and “Gregory Lemarchal” to KM. We thank Kenneth W. Bayles from the University of Nebraska Medical Center (Omaha) for providing *S. aureus* UAMS-1 WT and mutant strains.

## Conflict of interest

All the authors declare no competing interests.

## Ethical statement

All the strains used in this study were collected as part of the periodic monitoring of patients at the Hospices Civils de Lyon. This study was submitted to the Ethics Committee of the Hospices Civils de Lyon (HCL) and registered under CNIL No 17-216. All the patients were informed of the study; however, as the study was non-interventional no written informed consent was required under local regulations.

## Notes

### Competing Interest Statement

The authors have declared no competing interest.

