## Supplementary data for "Trophic cooperation promotes bacterial survival of *Staphylococcus aureus* and *Pseudomonas aeruginosa*"

**Materials and methods:** Generation of PA2600 knock-out Δ*acoR* and Δ*aco* mutants.

**Table S1:** Clinical CF strains used in this study.
**Table S2:** Non-CF strains and plasmids used in this study.
**Table S3:** Primers used in this study.
**Table S4:** List of *P. aeruginosa* genes differentially expressed in presence of *S. aureus* in the context of a competitive interaction.
**Table S5:** List of *P. aeruginosa* genes differentially expressed in presence of *S. aureus* in the context of coexistence.

**Figure S1:** Schematic representation of the employed methodology.
**Figure S2:** *P. aeruginosa* metabolic pathways and associated genes up-regulated or down-regulated in coexistence with *S. aureus*.
**Figure S3:** Acetoin concentration in supernatant of *S. aureus*, *B. subtilis*, *B. cenocepacia* and *S. maltophilia* monocultures or co-cultures with *P. aeruginosa*.
**Figure S4:** Fold change of *P. aeruginosa acoR*, *PA4148*, *liuA* and *zwf* gene expression induced by culture with *S. aureus* or its supernatant.
**Figure S5:** Monitoring of acetoin concentration in *S. aureus* and *P. aeruginosa* monocultures or co-culture or in *S. aureus* supernatant inoculated with *P. aeruginosa*, for the pairs SA146/PA146 and SA153/PA153A.
**Figure S6:** Acetoin concentration in CF sputa from patients.
**Figure S7:** Acetoin concentration in cultures of *S. aureus* strains from competition and coexistence couples.
**Figure S8:** Glucose concentrations in cultures of *S. aureus* and *P. aeruginosa* strains from competition and coexistence pairs.
**Figure S9:** Growth kinetic of *S. aureus* cultivated in absence or presence of acetoin.

### Materials and methods

**Generation of PA2600 knock-out Δ*acoR* and Δ*aco* mutants:** upstream and downstream flanking regions of *acoR* and *aco* operon (474 bp and 486 bp fragments for *acoR* ; 654 bp and 708 bp for *aco*) were PCR amplified (GoTaq polymerase, Promega) and cloned into pEXG2 by Sequence Ligation and Independent Cloning (SLIC) method (1,2). Resulting plasmids pEXG2-*acoR* and pEXG2-*aco* were then transferred into PA2600 by triparental mating. A first conjugation was performed between the two *E. coli* strains carrying either the pRK2013 helper plasmid or the constructed pEXG2 plasmid by spotting 30µL of pre-culture of each strain on LB plates. After two hours at 37°C, 30µL of PA2600 pre-culture were added on the dried spot and the plate was incubated five hours at 37°C. The spot was then re-suspended in LB medium and plated on Cetrimide plates supplemented with gentamycin. Resulting clones were then plated on LB containing 10% sucrose to select for plasmid excision by crossing-over. The resulting strains were checked for gentamicin sensitivity and gene deletion by PCR.

| CF clinical strains | CF patient | Strain name | Interaction state | Experiments | Reference |
|---|---|---|---|---|---|
| *P. aeruginosa* (PA) and *S. aureus* (SA) | 1 | PA2596 | Competition (SA2599, SA2597) | RNAseq | 3 |
| | | SA2597 | | RNAseq | 3 |
| | 2 (■) | PA2600 | Coexistence (SA2599, SA2597) | RNAseq; qRT-PCR screening; *aco* induction qRT-PCR; Acetoin monitoring in co-culture; Acetoin catabolism in SA supernatant; Culture in minimal medium; 5-day co-cultures | 3 |
| | | SA2599 | | RNA seq; qRT-PCR screening; *aco* induction qRT-PCR; Acetoin production screening; Acetoin monitoring in co-culture; Production of SA supernatant; 5 days co-cultures; Growth in presence of acetoin | 3 |
| | 3 | PA7A | Coexistence | qRT-PCR screening | This study |
| | | SA7 | | qRT-PCR screening | This study |
| | 4 | PA13 | Coexistence | qRT-PCR screening | This study |
| | | SA13 | | qRT-PCR screening | This study |
| | 5 (●) | PA27 | Coexistence | qRT-PCR screening | 3 |
| | | SA27 | | qRT-PCR screening | 3 |
| | 6 | PA30 | Coexistence | qRT-PCR screening | 3 |
| | | SA30 | | qRT-PCR screening | 3 |
| | 7 | PA31 | Coexistence | qRT-PCR screening | 3 |
| | | SA31 | | qRT-PCR screening | 3 |
| | 8 | PA37 | Coexistence | qRT-PCR screening | This study |
| | | SA37 | | qRT-PCR screening | This study |
| | 9 | PA42 | Coexistence | qRT-PCR screening | 3 |
| | | SA42 | | qRT-PCR screening | 3 |
| | 10 | PA48 | Coexistence | qRT-PCR screening | This study |
| | | SA48 | | qRT-PCR screening | This study |
| | 11 | PA53 | Coexistence | qRT-PCR screening | This study |
| | | SA53 | | qRT-PCR screening | This study |
| | 12 | PA54 | Coexistence | qRT-PCR screening | This study |
| | | SA54 | | qRT-PCR screening | This study |
| | 13 | PA69 | Coexistence | qRT-PCR screening | 3 |
| | | SA69 | | qRT-PCR screening | 3 |
| | 14 (▲) | PA80 | Coexistence | qRT-PCR screening | 3 |
| | | SA80 | | qRT-PCR screening | 3 |
| | 15 | PA82 | Coexistence | qRT-PCR screening | 3 |
| | | SA82 | | qRT-PCR screening | 3 |
| | 16 | PA146 | Coexistence | qRT-PCR screening; Acetoin catabolism screening; Acetoin monitoring in co-culture; Acetoin catabolism in SA supernatant | 3 |
| | | SA146 | | qRT-PCR screening; Acetoin production screening; Acetoin monitoring in co-culture; Production of SA supernatant | 3 |
| | 14 (▲) | PA148B | Coexistence | qRT-PCR screening; Acetoin catabolism screening | 3 |
| | | SA148 | | qRT-PCR screening; Acetoin production screening | 3 |
| | 17 | PA152 | Coexistence | qRT-PCR screening; Acetoin catabolism screening | 3 |
| | | SA152 | | qRT-PCR screening; Acetoin production screening | 3 |
| | 18 | PA153A | Coexistence | qRT-PCR screening; Acetoin catabolism screening; Acetoin monitoring in co-culture; Acetoin catabolism in SA supernatant | 3 |
| | | PA153B | | Transcriptomic (qRT-PCR) | 3 |
| | | SA153 | | qRT-PCR screening; Acetoin production screening; Acetoin monitoring in co-culture; Production of SA supernatant | 3 |

| | | | | | |
|---|---|---|---|---|---|
| | 19 | PA154A | Coexistence | qRT-PCR screening; Acetoin catabolism screening | 3 |
| | | PA154B | | qRT-PCR screening; Acetoin catabolism screening | 3 |
| | | SA154 | | qRT-PCR screening; Acetoin production screening | 3 |
| | 20 | PA156 | Coexistence | qRT-PCR screening; Acetoin catabolism screening | 3 |
| | | SA156 | | qRT-PCR screening; Acetoin production screening | 3 |
| | 21 | PA166A | Coexistence | qRT-PCR screening; Acetoin catabolism screening | 3 |
| | | SA166 | | qRT-PCR screening; Acetoin production screening | 3 |
| | 22 | PA167 | Competition | Acetoin catabolism screening | 3 |
| | | SA167 | | Acetoin production screening | 3 |
| | 2 (■) | PA171A | Coexistence | Acetoin catabolism screening | 3 |
| | | SA171 | | Acetoin production screening | 3 |
| | 23 | PA172 | Competition | Acetoin catabolism screening | This study |
| | | SA172 | | Acetoin production screening | This study |
| | 24 | PA178 | Coexistence | Acetoin catabolism screening | 3 |
| | | SA178 | | Acetoin production screening | 3 |
| | 25 | PA179 | Competition | Acetoin catabolism screening | 3 |
| | | SA179 | | Acetoin production screening | 3 |
| | 26 | PA181 | Competition | Acetoin catabolism screening | 3 |
| | | SA181 | | Acetoin production screening | 3 |
| | 27 (♦) | PA186 | Competition | Acetoin catabolism screening | 3 |
| | | SA186 | | Acetoin production screening | 3 |
| | 28 | PA187 | Competition | Acetoin catabolism screening | This study |
| | 29 | PA188A | Competition | Acetoin catabolism screening | This study |
| | | SA188 | | Acetoin production screening | This study |
| | 30 | PA193A | Competition | Acetoin catabolism screening | This study |
| | | SA193 | | Acetoin production screening | This study |
| | 5 (●) | PA194A | Coexistence | Acetoin catabolism screening | This study |
| | | PA194B | | Acetoin catabolism screening | This study |
| | | SA194 | | Acetoin production screening | This study |
| | 31 | PA197 | Competition | Acetoin catabolism screening | This study |
| | | SA197 | | Acetoin production screening | This study |
| | 32 | SA198 | Competition | Acetoin production screening | This study |
| | 33 | PA199A | Competition | Acetoin catabolism screening | This study |
| | | PA199C | | Acetoin catabolism screening | This study |
| | 34 | PA200 | Competition | Acetoin catabolism screening | This study |
| | | SA200 | | Acetoin production screening | This study |
| | 35 | SA205 | Competition | Acetoin production screening | This study |
| | 36 | SA207 | Coexistence | Acetoin production screening | This study |
| | 27 (♦) | SA213 | Competition | Acetoin production screening | This study |
| *B. cenocepacia* | 37 | LUG2886 | Coexistence (PA2600) | qRT-PCR (*aco* induction) | This study |
| *S. maltophilia* | 38 | LUG2884 | Coexistence (PA2600) | qRT-PCR (*aco* induction) | This study |

**Table S1: Clinical CF strains used in this study.** Unless indicated, interaction state was tested between strains from the same clinical pair (*ie*. isolated from a single patient). Four strain pairs were recovered from a same patient but at different time points and are annotated with identical patient number and symbol (■,●,▲ or ♦).

| Strains / plasmids | Name | Characteristics | Interaction state | Experiments | Reference |
|---|---|---|---|---|---|
| *P. aeruginosa* | PA2600 Δ*acoR* | *acoR* deletion mutant | Coexistence (SA2599) | Culture in minimal medium; 5-day co-cultures | This study |
| | PA2600 Δ*aco* | *aco* operon deletion mutant | Coexistence (SA2599) | Culture in minimal medium; 5-day co-cultures | This study |
| *S. aureus* | SA UAMS-1 | WT strain | Coexistence (PA2600) | Production of SA supernatant (*aco* induction) | 4 |
| | SA UAMS-1 Δ*alsSD* | UAMS-1489, *alsSD* deletion mutant | Coexistence (PA2600) | Production of SA supernatant (*aco* induction) | 4 |
| *B. subtilis* | LUG2953 | WT strain | Coexistence (PA2600) | qRT-PCR (*aco* induction) | This study |
| Plasmids | pEXG2 | $Gm^R$ ; mobilizable, non-replicative vector in *P. aeruginosa* | | *acoR* and *aco* deletions | 2 |
| | pEXG2-ΔacoR | pEXG2 carrying upstream and downstream sequences of *acoR* for gene deletion | | *acoR* deletion | This study |
| | pEXG2-Δaco | pEXG2 carrying upstream and downstream sequences of *aco* for operon deletion | | *aco* deletion | This study |
| | pRK2013 | $Km^R$, helper plasmid with conjugative properties | | *acoR* and *aco* deletions | 5 |

**Table S2: Non-CF strains and plasmids used in this study.** *P. aeruginosa* Δ*acoR* and Δ*aco* mutants were constructed from the clinical CF isolate PA2600 (Table S1). Interaction state was tested with the strain indicated in brackets.

| Use | Name | Sequence | Target | Amplicon Size |
|---|---|---|---|---|
| qPCR | PArpoD-F | GCGCAACAGCAATCTCGTCT | *rpoD* | 177 |
| | PArpoD-R | ATCCGGGGCTGTCTCGAATA | | |
| | PAgyrB-F | ATCTCGGTGAAGGTACCGGA | *gyrB* | 160 |
| | PAgyrB-R | TGCCTTCGTTGGGATTCTCC | | |
| | OLC8-F | GCGAGGATCTCTACTTCCGC | *acoR* | 140 |
| | OLC8-R | CTCACCGAGTTCGATGCGTA | | |
| | OLC9-F | GCGGATCGTCAACCTGTCAT | *PA4148* | 86 |
| | OLC9-R | CGATCACGGCAAACTTCGAG | | |
| | OLC14-F | GTTCTTCAGGCTCCAGTCGG | *pvdS* | 81 |
| | OLC14-R | TTGCGGACGATCTGGAACAG | | |
| | OLC16-F | CGCGACAAGAGCGAATACCT | *lasA* | 71 |
| | OLC16-R | AGGGTCAGCAACACTTTCGG | | |
| | OLC17-F | CTGGACTGAACCAGGCGATG | *rhlA* | 113 |
| | OLC17-R | CAGGTATTTGCCGACGGTCT | | |
| | OLC22-F | CGACGGTATCCAGGTCGATG | *PA1874* | 70 |
| | OLC22-R | GAACTTGACCGTGACCACCT | | |
| | OLC23-F | CGGTGTTGCTCGGATACCTC | *rcpC* | 120 |
| | OLC23-R | GCTTGCGTTCGAGCTTTTCC | | |
| | OLC25-F | ATCGCCTGCTTCCAGTTGTC | *pchD* | 166 |
| | OLC25-R | AGAGAGTGAAGTTGTGCGCC | | |
| | OLC29-F | GAGCGACGAACTGACCTACC | *rhlB* | 200 |
| | OLC29-R | TACTTCTCGTGAGCGATGCG | | |
| | OLC31-F | TGCTCACTTCGCTATGGACC | *acoB* | 117 |
| | OLC31-R | AGAAGATCACCGGGTCGTTG | | |
| | OLC33-F | CCGACGTGATCGCCTTCATA | *PA4153* | 83 |
| | OLC33-R | GACGATTTCTTCCAGGCCGA | | |
| | OLC35-F | GCCCATCGAGTTCCGTATGT | *bauD* | 151 |
| | OLC35-R | GAGGCAGACGGTGAAGATCG | | |
| | OLC39-F | GACGAAGACGGCATGAACCT | *tadA* | 119 |
| | OLC39-R | TCCAGTTCGTAGCGGGAGAT | | |
| | OLC41-F | CCACCACGAAACTGCAACTG | *tadG* | 107 |
| | OLC41-R | GCTTCTCCAACCGAAGTCCA | | |
| | OLC54-F | TTGCCGTATTGAGTCCCACG | *flp* | 77 |
| | OLC54-R | GACTTTTTCGCCGACTCCGT | | |
| | OLC56-F | GGACTGACGCTCAGGCAAT | *rhlC* | 74 |
| | OLC56-R | CCGGAGGAGATCAGGAACGA | | |
| | OLC71-F | CAGCCGGACGAAGGTATCTC | *zwf* | 113 |
| | OLC71-R | GCGTGGTAGGTCTCGGAAAA | | |
| | OLC72-F | GCTATACCCGCTCAAGGGC | *dguA* | 92 |
| | OLC72-R | TCTTGCGGTCGTAATCGGTG | | |
| | OLC73-F | CTTCGCCTACCTGTTCAGCC | *PA5099* | 163 |
| | OLC73-R | CGGCAGGTAGCGTGAATAGT | | |
| | OLC74-F | GCAGTTCGGAGCACATCAAC | *PA1874* | 130 |
| | OLC74-R | TCGATGCTGACGAAATCGGT | | |
| | OLC75-F | TGGTATCCGGCGAACACATC | *liuA* | 127 |
| | OLC75-R | CACATCTTGCTGCCGTTGAG | | |
| | OLC92-F | CAGCCAGGACTACGAGAACG | *lasR* | 153 |

| | | | | |
|---|---|---|---|---|
| | OLC92-R | TGGTAGATGGACGGTTCCCA | | |
| | OLC93-F | GGCATTCCCCTCACCGAC | *gntK* | 181 |
| | OLC93-R | GGGTCAGTTCCAGGTAGACGA | | |
| | OLC94-F | TGGAAGTGGCTGCTCAATCC | *gltF* | 81 |
| | OLC94-R | CCAGTCGAAGCGGAAACCTT | | |
| | OLC95-F | CACCTGCACCTTCTATGGCA | *edd* | 96 |
| | OLC95-R | GTGTTCGGGTTGACGAAGGA | | |
| | OLC96-F | CAACGGCACCTCGATCTTCA | *mmsA* | 94 |
| | OLC96-R | AATCGGGATGTTGATGCCCA | | |
| Cloning | OLC58-F | GGTCGACTCTAGAGGATCCCC AGGGCGATGCCCCGGCCGATG | *acoR* downstream | 474 |
| | OLC58-R | CACATGGTCCTTCGAGTGTGC | | |
| | OLC59-F | GCACACTCGAAGGACCATGTG CGCCGGCATGGCATCCGCATG | *acoR* upstream | 486 |
| | OLC59-R | ACCGAATTCGAGCTCGAGCCC CTCGATCGCGCGGACGAACCA | | |
| | OLC64-F | GGTCGACTCTAGAGGATCCCC TAGTCGGCATCGCGCACCCGT | *aco* operon downstream | 654 |
| | OLC64-R | GGAGAAATCGTCGGACGACGT | | |
| | OLC65-F | ACGTCGTCCGACGATTTCTCCA GTTGGTGAACAACAAGGAGC | *aco* operon upstream | 708 |
| | OLC65-R | ACCGAATTCGAGCTCGAGCCC ACAGCCTGACCACTTTCGTGC | | |

**Table S3: Primers used in this study.**

| Gene (clinical strains) | Gene (PAO1) | | | UniProt | Product | Function | SA2599/PA2596 | | SA2597/PA2596 | | LasR regulation | Ref. |
|---|---|---|---|---|---|---|---|---|---|---|---|---|
| | | | | | | | $Log_2$ fold-change | Adjusted p-value | $Log_2$ fold-change | Adjusted p-value | | |
| fecI_2 | **NA** | **NA** | | P23484 | Putative RNA polymerase sigma factor FecI | Iron metabolism | **-2.67** | 4.02E-06 | **-2.51** | 1.85E-04 | | 6 |
| group_1318 | **NA** | **NA** | | P40883 | Regulatory protein PchR | Iron metabolism | **-3.57** | 3.88E-23 | **-3.13** | 6.04E-25 | | 6 |
| group_1539 | **NA** | **NA** | | NA | Hypothetical protein | Unknown | **-2.96** | 3.51E-11 | **-3.58** | 3.20E-13 | | |
| bauA_1 | ***PA0132*** | ***bauA*** | | Q9I700 | Beta-alanine-pyruvate aminotransferase | Aminoacids (ILV) catabolism, propanoate metabolism | **3.15** | 1.00E-06 | **3.44** | 6.25E-39 | Activation | 7 |
| nirS_1 | ***PA0509*** | ***nirN*** | a | Q9I609 | Nitrite reductase | Nitrogen metabolism | **-2.92** | 6.77E-10 | **-2.93** | 3.30E-09 | Repression | 8,9 |
| cysG_2 | ***PA0510*** | ***nirE*** | a | G3XD80 | Siroheme synthase NirE | Nitrogen, porphyrin and chlorophyll metabolism | **-3.00** | 8.64E-09 | **-2.56** | 5.61E-06 | Repression | 8,9 |
| group_5012 | ***PA0512*** | ***nirH*** | a | P95415 | NirH | Nitrogen metabolism | **-3.27** | 8.64E-09 | **-2.23** | 4.94E-04 | Repression | 8,9 |
| group_3026 | ***PA0513*** | ***nirG*** | a | P95414 | NirG | Nitrogen metabolism | **-2.79** | 5.29E-06 | **-2.35** | 9.60E-04 | | 8 |
| group_4848 | ***PA0514*** | ***nirL*** | a | P95413 | Heme d1 biosynthesis protein NirL | Nitrogen metabolism | **-3.09** | 1.71E-08 | **-3.47** | 2.61E-08 | | 8 |
| group_3339 | ***PA0515*** | ***nirD*** | a | P95412 | Probable transcriptional regulator | Nitrogen metabolism | **-2.99** | 2.04E-07 | **-2.03** | 1.30E-03 | | 8 |
| group_5595 | ***PA0516*** | ***nirF*** | a | Q51480 | Heme d1 biosynthesis protein NirF | Nitrogen metabolism | **-3.34** | 1.03E-10 | **-3.47** | 6.65E-16 | | 8 |
| nirS_2 | ***PA0519*** | ***nirS*** | a | P24474 | Nitrite reductase | Nitrogen metabolism | **-3.29** | 3.53E-07 | **-5.39** | 6.48E-51 | | 8 |
| nirQ | ***PA0520*** | ***nirQ*** | b | Q51481 | Denitrification regulatory protein NirQ | Nitrogen metabolism | **-4.27** | 3.41E-28 | **-4.51** | 3.08E-33 | | 8 |
| qoxC | ***PA0521*** | ***nirO*** | b | G3XD44 | Quinol oxidase subunit 3 | Unknown | **-4.65** | 2.60E-22 | **-4.60** | 4.38E-15 | | |
| group_6044 | ***PA0522*** | ***nirP*** | b | Q51483 | Hypothetical protein | Unknown | **-2.90** | 1.59E-05 | **-2.99** | 7.07E-05 | | |
| norB | ***PA0524*** | ***norB*** | | Q59647 | Nitric oxide reductase subunit B | Nitrogen metabolism | **-5.53** | 1.99E-23 | **-8.08** | 1.91E-79 | | 8 |
| group_3360 | ***PA0672*** | ***hemO*** | | G3XCZ8 | Heme oxygenase | Porphyrin and chlorophyll metabolism | **-2.38** | 9.28E-05 | **-2.53** | 1.57E-17 | | |
| ppsC | ***PA1137*** | ***PA1137*** | | Q9I4J8 | Phthiocerol/phenolphthiocerol synthesis polyketide synthase | Unknown | **-4.31** | 1.74E-21 | **-3.55** | 1.62E-19 | | |
| rocR | ***PA1196*** | ***ddaR*** | | Q9I4E2 | Arginine utilization regulatory protein RocR | Aminoacids (arginine) metabolism | **2.51** | 3.23E-05 | **3.22** | 5.81E-21 | | 10 |
| fecI_5 | ***PA1300*** | ***PA1300*** | c | Q9I444 | Putative RNA polymerase sigma factor FecI | Iron metabolism | **-3.17** | 3.51E-11 | **-2.50** | 2.25E-11 | | 6 |
| fecR_1 | ***PA1301*** | ***PA1301*** | c | Q9I443 | Protein FecR | Iron metabolism | **-3.02** | 1.71E-08 | **-2.53** | 1.01E-13 | | 6 |
| ctpF | ***PA1429*** | ***PA1429*** | | Q9I3R5 | Putative cation-transporting ATPase F | Unknown | **2.89** | 8.45E-09 | **3.63** | 7.15E-35 | | |
| group_3703 | ***PA1673*** | ***PA1673*** | | Q9I352 | Bacteriohemerythrin | Unknown | **2.16** | 1.57E-04 | **2.57** | 1.30E-14 | | |
| group_5526 | ***PA1746*** | ***PA1746*** | | Q9I2Z1 | Hypothetical protein | Unknown | **2.42** | 1.59E-06 | **2.42** | 1.38E-09 | | |
| group_2540 | ***PA1747*** | ***PA1747*** | | Q9I2Z0 | Hypothetical protein | Unknown | **2.57** | 8.64E-09 | **2.46** | 1.07E-08 | | |
| group_269 | ***PA2033*** | ***PA2033*** | | Q9I282 | Hypothetical protein | Unknown | **-2.24** | 1.65E-03 | **-2.84** | 7.11E-23 | | |
| ybdM | ***PA2126*** | ***cgrC*** | | Q9I1Y9 | CupA gene regulator C, CgrC | Unknown | **2.60** | 3.18E-07 | **2.46** | 1.05E-06 | | |
| group_6509 | ***PA2321*** | ***gntK*** | | G3XD53 | Gluconokinase | Carbon metabolism (pentose phosphate) | **-2.73** | 4.03E-06 | **-2.41** | 2.68E-05 | Repression | 9 |
| group_3203 | ***PA2384*** | ***PA2384*** | | Q9I195 | Ferric uptake regulation protein | Iron metabolism | **-2.44** | 1.09E-04 | **-2.56** | 8.25E-10 | | 6 |
| mbtH | ***PA2412*** | ***PA2412*** | | Q9I169 | Hypothetical protein | Monobactam biosynthesis | **-2.12** | 1.66E-03 | **-2.27** | 6.19E-08 | | |
| group_3129 | ***PA2468*** | ***foxI*** | | Q9I114 | Putative RNA polymerase sigma factor FecI | Iron metabolism | **-2.32** | 1.63E-07 | **-2.03** | 1.07E-08 | | 6 |
| hmp | ***PA2664*** | ***fhp*** | | Q9I0H4 | Flavohemoprotein | Iron metabolism, NO detoxification | **-2.41** | 1.85E-05 | **-2.38** | 1.86E-03 | | 11 |
| group_3982 | ***PA2691*** | ***PA2691*** | | Q9I0F1 | NADH dehydrogenase-like protein | Oxidative phosphorylation | **-4.38** | 1.03E-18 | **-3.45** | 3.32E-07 | | |
| hldD | ***PA3337*** | ***rfaD*** | | Q9HYQ8 | ADP-L-glycero-D-manno-heptose-6-epimerase | Lipopolysaccharide biosynthesis | **3.40** | 1.45E-12 | **4.26** | 1.62E-48 | | |
| yccM | ***PA3391*** | ***nosR*** | d | Q9HYL3 | Regulatory protein NosR | Nitrogen metabolism | **-3.57** | 7.09E-14 | **-4.08** | 1.98E-30 | Repression | 8,9 |
| nosZ | ***PA3392*** | ***nosZ*** | d | Q9HYL2 | Nitrous-oxide reductase | Nitrogen metabolism | **-3.62** | 1.57E-10 | **-5.50** | 6.62E-51 | Repression | 8,9 |
| nosD | ***PA3393*** | ***nosD*** | d | Q9HYL1 | Putative ABC transporter binding protein NosD | Nitrogen metabolism | **-3.13** | 1.38E-10 | **-3.97** | 3.44E-20 | Repression | 8,9 |
| nosY | ***PA3395*** | ***nosY*** | d | Q9HYK9 | Putative ABC transporter permease protein NosY | ABC transporters (nitrogen metabolism) | **-3.14** | 1.69E-07 | **-2.91** | 1.48E-06 | Repression | 8,9 |

| | | | | | | | | | | | | |
|---|---|---|---|---|---|---|---|---|---|---|---|---|
| nosL | ***PA3396*** | ***nosL*** | d | Q9HYK8 | Copper-binding lipoprotein NosL | Nitrogen metabolism | **-2.60** | 3.86E-05 | **-2.46** | 5.00E-05 | Repression | 8,9 |
| group_3537 | ***PA3411*** | ***PA3411*** | | Q9HYJ4 | Hypothetical protein | Unknown | **-2.59** | 3.63E-06 | **-2.05** | 1.41E-03 | | |
| bfd | ***PA3530*** | ***bfd*** | | Q9HY80 | Bacterioferritin-associated ferredoxin | Unknown | **-2.24** | 3.28E-10 | **-2.04** | 1.81E-10 | | |
| ykgO | ***PA3600*** | ***rpl36*** | e | Q9HY26 | 50S ribosomal protein L36 2 | Ribosome structure | **-2.99** | 6.77E-10 | **-2.87** | 3.60E-08 | | |
| rpmE2 | ***PA3601*** | ***ykgM*** | e | Q9HY25 | 50S ribosomal protein L31 type B | Ribosome structure | **-2.92** | 2.37E-10 | **-3.21** | 2.69E-26 | | |
| fdx_1 | ***PA3809*** | ***fdx2*** | f | Q51383 | 2Fe-2S ferredoxin | Iron-sulfur protein | **-2.72** | 4.17E-12 | **-3.02** | 7.98E-16 | | |
| hscA | ***PA3810*** | ***hscA*** | f | Q51382 | Chaperone protein HscA | Protein stabilization | **-2.56** | 4.41E-11 | **-2.60** | 6.55E-16 | | 12 |
| hscB | ***PA3811*** | ***hscB*** | f | Q9HXJ1 | Co-chaperone protein HscB | Protein stabilization | **-2.04** | 1.40E-06 | **-2.28** | 1.35E-14 | | 12 |
| iscA | ***PA3812*** | ***iscA*** | f | Q9HXJ0 | Iron-binding protein IscA | [Fe-S] cluster biogenesis | **-2.51** | 2.80E-10 | **-3.37** | 7.39E-26 | | 13 |
| iscU | ***PA3813*** | ***iscU*** | f | Q9HXI9 | Iron-sulfur cluster assembly scaffold protein IscU | [Fe-S] cluster biogenesis | **-2.40** | 3.46E-06 | **-3.34** | 1.95E-27 | | 13 |
| iscS_1 | ***PA3814*** | ***iscS*** | f | Q9HXI8 | Cysteine desulfurase IscS | Sulfur relay system, thiamine metabolism, [Fe-S] cluster biogenesis | **-2.24** | 5.79E-05 | **-3.44** | 3.85E-29 | | 13 |
| iscR | ***PA3815*** | ***iscR*** | f | Q9HXI7 | HTH-type transcriptional regulator IscR | [Fe-S] cluster biogenesis regulation | **-2.42** | 4.44E-06 | **-3.65** | 4.38E-31 | | 13 |
| copA_4 | ***PA3920*** | ***yvgX*** | | Q9HX93 | Copper-exporting P-type ATPase | Unknown | **-2.28** | 3.75E-07 | **-2.63** | 4.15E-14 | | |
| ntaA | ***PA4155*** | ***PA4155*** | | Q9HWM6 | Nitrilotriacetate monooxygenase component A | Unknown | **-3.09** | 5.86E-10 | **-2.23** | 6.24E-06 | | |
| fyuA_1 | ***PA4156*** | ***fvbA*** | | Q9HWM5 | Pesticin receptor | Iron metabolism | **-2.70** | 8.32E-12 | **-2.48** | 2.30E-09 | | 14 |
| fepC_1 | ***PA4158*** | ***fepC*** | | Q9HWM3 | Ferric enterobactin transport ATP-binding protein FepC | ABC transporters (iron complex) | **-2.30** | 6.59E-05 | **-2.21** | 3.50E-04 | | |
| fepB | ***PA4159*** | ***fepB*** | | Q9HWM2 | Ferrienterobactin-binding periplasmic protein | ABC transporters (iron complex) | **-3.21** | 1.20E-10 | **-2.48** | 7.35E-06 | | |
| zupT | ***PA4467*** | ***PA4467*** | g | Q9HVV1 | Zinc transporter ZupT | Unknown | **-3.12** | 1.92E-08 | **-2.97** | 1.40E-15 | | |
| sodB_2 | ***PA4468*** | ***sodM*** | g | P53652 | Superoxide dismutase [Mn/Fe] | Oxydative and iron stress response | **-3.17** | 1.19E-09 | **-3.09** | 4.37E-19 | | 15 |
| group_2514 | ***PA4469*** | ***PA4469*** | g | Q9HVV0 | Hypothetical protein | Unknown | **-3.32** | 8.79E-09 | **-3.28** | 2.91E-23 | | |
| fumC_1 | ***PA4470*** | ***fumC1*** | g | Q51404 | Fumarate hydratase class II | Carbon metabolism (cytrate circle), iron stress response | **-3.14** | 6.08E-08 | **-3.16** | 2.75E-18 | | 15 |
| group_4225 | ***PA4471*** | ***fagA*** | g | G3XD99 | Hypothetical protein | Unknown | **-3.52** | 3.14E-17 | **-3.23** | 1.64E-25 | | |
| group_3398 | ***PA4570*** | ***PA4570*** | | Q9HVL4 | Hypothetical protein | Unknown | **-4.12** | 3.14E-17 | **-4.32** | 4.87E-31 | | |
| ccpA | ***PA4587*** | ***ccpR*** | | P14532 | Cytochrome c551 peroxidase | Oxydative stress response | **3.16** | 3.49E-11 | **3.27** | 6.90E-16 | Repression | 7,16 |
| group_1579 | ***PA4625*** | ***cdrA*** | | Q9HVG6 | Cyclic diguanylate-regulated TPS partner A, CdrA | Adhesion and biofilm matrix structure | **2.53** | 8.83E-05 | **2.06** | 6.78E-10 | | 17 |
| fecI_3 | ***PA4896*** | ***PA4896*** | | Q9HUR7 | Putative RNA polymerase sigma factor FecI | Iron metabolism | **-3.04** | 6.78E-10 | **-2.23** | 1.49E-08 | | 6 |
| group_4196 | ***PA5027*** | ***PA5027*** | | Q9HUE2 | Hypothetical protein | Unknown | **2.35** | 7.70E-05 | **2.82** | 2.70E-20 | Activation | 9 |
| arcD | ***PA5170*** | ***arcD*** | | P18275 | Arginine/ornithine antiporter | Aminoacids (arginine) metabolism | **2.36** | 4.52E-04 | **2.49** | 5.13E-04 | | 10 |
| NA | ***PA5369.3*** | ***PA5369.3*** | | NA | tRNA-Ala | Aminoacyl-tRNA biosynthesis | **2.12** | 7.09E-03 | **4.50** | 3.69E-21 | | |
| pka | ***PA5475*** | ***PA5475*** | | Q9HT95 | Protein lysine acetyltransferase Pka | Unknown | **2.04** | 4.85E-04 | **2.30** | 1.81E-10 | | |

**Table S4: List of *P. aeruginosa* genes differentially expressed in presence of *S. aureus* in the context of a competitive interaction.** PA2596 competition strain was cultivated in absence or presence of SA2599 or SA2597. RNAs were extracted after 4 hours of culture and RNAseq analysis was performed as described in material and methods. A gene was considered as differentially expressed when the Fold Change (FC) was > $|2\log_2|$ with an adjusted *P*-value<0.05 in presence of both SA strains. Genes from the same operon are annotated with an identical letter. Grey cells indicate genes that were also dysregulated in the context of coexistence (Table S5). Functional classification was performed thanks to KEGG database and literature.

| Gene (clinical strains) | Gene (PAO1) | | | UniProt | Product | Function | SA2599/PA2600 | | SA2597/PA2600 | | LasR regulation | Ref. |
|---|---|---|---|---|---|---|---|---|---|---|---|---|
| | | | | | | | $Log_2$ fold-change | Adjusted p-value | $Log_2$ fold-change | Adjusted p-value | | |
| group_955 | **NA** | **NA** | | NA | Hypothetical protein | Unknown | **-2,20** | 3,84E-09 | **-2,56** | 1,54E-05 | | |
| eamB_1 | **NA** | **NA** | | P38101 | Cysteine/O-acetylserine efflux protein | Aminoacids transport | **2,04** | 5,37E-04 | **2,31** | 2,69E-03 | | |
| group_511 | **NA** | **NA** | | NA | Hypothetical protein | Unknown | **-2,04** | 2,34E-03 | **-2,77** | 5,28E-03 | | |
| ▶ bauD | ***PA0129*** | ***bauD*** | | Q9I703 | Putative GABA permease | Aminoacids (β-alanine) catabolism | **2,90** | 1,53E-15 | **3,07** | 2,83E-14 | | 18 |
| bauB | ***PA0131*** | ***bauB*** | | Q9I701 | Beta-alanine degradation protein BauB | Aminoacids (β-alanine) catabolism | **2,27** | 2,30E-05 | **2,61** | 2,31E-04 | | 18 |
| cysG_2 | ***PA0510*** | ***nirE*** | a | G3XD80 | Siroheme synthase NirE | Nitrogen, porphyrin and chlorophyll metabolism | **-2,42** | 6,73E-05 | **-2,72** | 1,36E-03 | Repression | 8,9 |
| group_3026 | ***PA0513*** | ***nirG*** | a | P95414 | NirG | Nitrogen metabolism | **-2,69** | 3,29E-05 | **-2,97** | 2,87E-03 | | 8 |
| group_3339 | ***PA0515*** | ***nirD*** | a | P95412 | Probable transcriptional regulator | Nitrogen metabolism | **-2,62** | 1,99E-03 | **-3,37** | 3,73E-04 | | 8 |
| group_5595 | ***PA0516*** | ***nirF*** | a | Q51480 | Heme d1 biosynthesis protein NirF | Nitrogen metabolism | **-3,53** | 1,60E-08 | **-3,88** | 1,44E-06 | | 8 |
| nirM | ***PA0518*** | ***nirM*** | a | P00099 | Cytochrome c-551 | Nitrogen metabolism | **-2,62** | 4,27E-03 | **-3,27** | 3,88E-03 | | 8 |
| nirS_2 | ***PA0519*** | ***nirS*** | a | P24474 | Nitrite reductase | Nitrogen metabolism | **-3,27** | 7,25E-06 | **-4,43** | 2,95E-08 | | 8 |
| nirQ | ***PA0520*** | ***nirQ*** | | Q51481 | Denitrification regulatory protein NirQ | Nitrogen metabolism | **-3,25** | 7,06E-05 | **-4,12** | 2,10E-06 | | 8 |
| norC | ***PA0523*** | ***norC*** | h | Q59646 | Nitric oxide reductase subunit C | Nitrogen metabolism | **-3,59** | 2,99E-05 | **-6,23** | 4,50E-14 | | 8 |
| norB | ***PA0524*** | ***norB*** | h | Q59647 | Nitric oxide reductase subunit B | Nitrogen metabolism | **-5,82** | 3,20E-18 | **-5,77** | 2,52E-12 | | 8 |
| group_4191 | ***PA0525*** | ***norD*** | h | Q51484 | Probable dinitrification protein NorD | Nitrogen metabolism | **-3,86** | 1,68E-06 | **-4,82** | 1,23E-08 | | 8 |
| group_364 | ***PA0526*** | ***PA0526*** | | G3XDA0 | Hypothetical protein | Unknown | **-3,38** | 6,53E-05 | **-4,60** | 3,45E-06 | | |
| NA | ***PA0668.3*** | **NA** | | NA | tRNA-Ala(tgc) | Aminoacyl-tRNA biosynthesis | **3,45** | 5,66E-05 | **7,57** | 1,86E-22 | | |
| acrC_2 | ***PA0746*** | ***PA0746*** | i | Q9I5I3 | Acryloyl-CoA reductase (NADH) | Unknown | **2,80** | 1,01E-12 | **3,12** | 2,02E-15 | | |
| mmsA_2 | ***PA0747*** | ***PA0747*** | i | Q9I5I2 | Methylmalonate-semialdehyde dehydrogenase | Aminoacids (isoleucine, leucine, valine) catabolism, propanoate and carbon metabolism | **2,08** | 1,04E-05 | **2,31** | 1,44E-06 | | |
| acnM | ***PA0794*** | ***PA0794*** | | Q9I5E4 | Aconitate hydratase A | Propanoate metabolism | **2,18** | 5,28E-10 | **2,90** | 1,80E-07 | | |
| aroP_2 | ***PA0866*** | ***aroP2*** | | Q9I575 | Aromatic amino acid transport protein AroP | Unknown | **2,30** | 8,73E-06 | **2,74** | 1,02E-03 | | |
| acs_1 | ***PA0887*** | ***acsA*** | | Q9I558 | Acetyl-coenzyme A synthetase | Propanoate, carbon and pyruvate metabolism | **3,08** | 5,58E-10 | **3,27** | 2,24E-07 | Repression | 7 |
| yybH | ***PA1325*** | ***yybH*** | j | Q9I419 | Putative protein YybH | Unknown | **2,16** | 1,03E-03 | **2,73** | 1,94E-05 | | |
| ilvA_1 | ***PA1326*** | ***ilvA*** | j | Q9I418 | L-threonine dehydratase biosynthetic IlvA | Carbon metabolism, aminoacids (isoleucine, leucine, valine) biosynthesis | **2,19** | 4,61E-05 | **2,46** | 6,93E-04 | | |
| ▶ group_1804 | ***PA1874*** | ***PA1874*** | k | Q9I2M3 | Hypothetical protein | Antibiotic resistance, quorum sensing | **-2,54** | 5,92E-05 | **-2,68** | 2,83E-04 | Activation | 9,19 |
| ▶ group_4895 | ***PA1875*** | ***PA1875*** | k | Q9I2M2 | Hypothetical protein | Antibiotic resistance | **-2,94** | 1,72E-09 | **-2,80** | 1,14E-05 | Activation | 9,19 |
| prsE_3 | ***PA1877*** | ***PA1877*** | k | Q9I2M0 | Type I secretion system membrane fusion protein PrsE | Antibiotic resistance | **-2,29** | 8,13E-03 | **-3,04** | 1,51E-07 | | 19 |
| group_6609 | ***PA1878*** | ***PA1878*** | | Q9I2L9 | Hypothetical protein | Unknown | **-2,03** | 2,47E-05 | **-2,38** | 1,01E-03 | | |
| group_2034 | ***PA1914*** | ***hvn*** | | Q9I2J0 | Hypothetical protein | Unknown | **-3,87** | 9,10E-16 | **-4,61** | 3,05E-20 | Activation | 7 |
| degU_4 | ***PA1978*** | ***erbR*** | | P29369 | Transcriptional regulatory protein DegU | Ethanol stress response | **2,28** | 2,43E-05 | **3,06** | 1,45E-07 | Repression | 9,20 |
| luxQ | ***PA1992*** | ***ercS*** | | Q9I2B7 | Autoinducer 2 sensor kinase/phosphatase LuxQ | Ethanol stress response | **2,13** | 9,50E-08 | **2,11** | 2,65E-04 | | 20 |
| accA1_2 | ***PA2012*** | ***liuD*** | l | Q9I299 | Acetyl-/propionyl-coenzyme A carboxylase alpha chain | Aminoacids (leucine) and monoterpenes catabolism | **2,30** | 7,81E-07 | **2,69** | 2,15E-06 | | 21 |
| menB | ***PA2013*** | ***liuC*** | l | Q9I298 | 1%2C4-dihydroxy-2-naphthoyl-CoA synthase | Aminoacids (leucine) and monoterpenes catabolism | **3,02** | 1,20E-11 | **3,29** | 7,09E-12 | | 21 |
| group_2911 | ***PA2014*** | ***liuB*** | l | Q9I297 | Methylmalonyl-CoA carboxyltransferase 12S subunit | Aminoacids (leucine) and monoterpenes catabolism | **2,61** | 1,20E-08 | **2,85** | 4,87E-10 | Activation | 9,21 |
| ▶ mmgC_7 | ***PA2015*** | ***liuA*** | l | Q9I296 | Acyl-CoA dehydrogenase | Aminoacids (leucine) and monoterpenes catabolism | **2,62** | 3,07E-08 | **2,76** | 5,26E-09 | | 21 |
| cueR_2 | ***PA2016*** | ***liuR*** | | Q9I295 | HTH-type transcriptional regulator CueR | Aminoacids (leucine) and monoterpenes catabolism | **2,61** | 4,02E-08 | **3,17** | 4,01E-06 | | 21 |

| | | | | | | | | | | | | | |
|---|---|---|---|---|---|---|---|---|---|---|---|---|---|
| | group_13 | *PA2040* | *pauA4* | | Q9I275 | Gamma-glutamylputrescine synthetase PuuA | Polyamines catabolism, aminoacids (glutamine) biosynthesis | 2,36 | 2,02E-08 | 2,77 | 8,62E-09 | | 22 |
| | kynU | *PA2080* | *kynU* | | Q9I235 | Kynureninase KynU | Aminoacids (tryptophan) catabolism | 2,63 | 9,29E-10 | 2,67 | 7,28E-07 | Activation | 7 |
| | group_1851 | *PA2166* | *PA2166* | | Q9I1U9 | Hypothetical protein | Unknown | -2,23 | 5,77E-05 | -2,82 | 1,56E-04 | Activation | 7 |
| | gntR_3 | *PA2320* | *gntR* | | Q9I1F6 | HTH-type transcriptional regulator GntR | Carbon metabolism (pentose phosphate) | -2,63 | 9,55E-11 | -2,71 | 1,51E-05 | | |
| ▶ | group_6509 | *PA2321* | *gntK* | | G3XD53 | Gluconokinase GntK | Carbon metabolism (pentose phosphate) | -3,67 | 1,18E-14 | -3,42 | 5,37E-06 | Repression | 9,23 |
| | group_1686 | *PA2462* | *PA2462* | | Q9I120 | Hypothetical protein | Unknown | 2,36 | 1,33E-10 | 2,34 | 7,08E-11 | | |
| | tsdA | *PA2481* | *PA2481* | m | Q9I101 | Thiosulfate dehydrogenase | Unknown | 2,68 | 1,47E-07 | 2,09 | 1,59E-04 | | |
| | group_3818 | *PA2482* | *PA2482* | m | Q9I100 | Cytochrome c4 | Unknown | 2,73 | 8,24E-08 | 2,39 | 6,15E-04 | | |
| | mmgC_5 | *PA2552* | *acdB* | n | Q9I0T2 | Acyl-CoA dehydrogenase | Unknown | 2,49 | 6,18E-12 | 2,82 | 2,35E-10 | Activation | 9 |
| | thlA_1 | *PA2553* | *PA2553* | n | Q9I0T1 | Acetyl-CoA acetyltransferase | Carbon and fatty acids metabolism | 2,46 | 7,14E-11 | 2,72 | 1,52E-08 | Activation | 9 |
| | group_5789 | *PA2554* | *PA2554* | n | Q9I0T0 | Putative oxidoreductase | Unknown | 2,78 | 1,75E-13 | 3,05 | 1,43E-09 | Activation | 9 |
| | acsA_1 | *PA2555* | *PA2555* | n | Q9I0S9 | Acetyl-coenzyme A synthetase | Propanoate, carbon and pyruvate metabolism | 2,83 | 1,51E-10 | 2,93 | 8,32E-07 | Activation | 9 |
| | fadD3 | *PA2557* | *PA2557* | | Q9I0S7 | 3-[(3aS%2C4S%2C7aS)-7a-methyl-1%2C5-dioxo-octahydro-1H-inden | Unknown | 3,04 | 2,30E-14 | 3,16 | 2,62E-11 | | |
| | lecA | *PA2570* | *lecA* | | Q05097 | PA-I galactophilic lectin LecA | Adhesion, biofilm formation | -3,40 | 1,59E-12 | -2,61 | 1,13E-03 | Activation | 7,9,24 |
| | group_5847 | *PA2662* | *PA2662* | o | Q9I0H6 | Hypothetical protein | Unknown | -3,18 | 1,66E-07 | -3,57 | 3,71E-05 | | |
| | group_3164 | *PA2663* | *ppyR* | o | Q9I0H5 | Psl and pyoverdine operon regulator, PpyR | Iron metabolism, biofilm formation and virulence | -2,83 | 7,91E-04 | -4,04 | 9,15E-07 | | 25 |
| | hmp | *PA2664* | *fhp* | | Q9I0H4 | Flavohemoprotein | Iron metabolism, NO detoxification | -5,40 | 1,82E-18 | -5,66 | 3,69E-14 | | 11 |
| | lip_2 | *PA2862* | *lipA* | | P26876 | Triacylglycerol lipase | Glycerolipid metabolism, virulence (lipase activity) | 2,31 | 6,63E-05 | 2,47 | 4,68E-04 | | |
| | group_2901 | *PA3038* | *opdQ* | | Q9HZH0 | Porin-like protein NicP | Membrane transports (in response to stress) | 2,75 | 8,42E-07 | 2,98 | 2,71E-06 | Repression | 7,26 |
| | ydfJ | *PA3079* | *PA3079* | p | Q9HZC9 | Membrane protein YdfJ | Unknown | 2,43 | 2,32E-07 | 2,29 | 2,55E-06 | | |
| | group_6150 | *PA3080* | *PA3080* | p | Q9HZC8 | Ycf48-like protein | Unknown | 2,33 | 5,77E-09 | 2,24 | 1,34E-04 | | |
| | eda_2 | *PA3181* | *edaA* | q | O68283 | 2-dehydro-3-deoxy-phosphogluconate aldolase | Carbon metabolism (pentose phosphate) | -2,85 | 2,30E-14 | -3,17 | 6,61E-11 | Activation | 7 |
| | pgl_1 | *PA3182* | *pgl* | q | Q9X2N2 | 6-phosphogluconolactonase | Carbon metabolism (pentose phosphate) | -3,10 | 1,45E-13 | -3,13 | 1,21E-08 | Activation | 7 |
| ▶ | zwf_2 | *PA3183* | *zwf* | q | O68282 | Glucose-6-phosphate 1-dehydrogenase | Carbon metabolism (pentose phosphate) | -2,77 | 4,39E-08 | -3,05 | 4,13E-11 | Activation | 7 |
| | ugpC | *PA3187* | *gltK* | r | Q9HZ51 | Sn-glycerol-3-phosphate import ATP-binding protein UgpC | ABC transporter (oligosaccharides, polyol, lipids and monosaccharides) | -3,88 | 1,55E-18 | -3,03 | 4,95E-08 | | |
| ▶ | ugpA | *PA3189* | *gltF* | r | Q9HZ49 | Sn-glycerol-3-phosphate transport system permease | ABC transporter (glucose/mannose) | -2,66 | 7,46E-08 | -2,25 | 4,99E-03 | Activation | 7 |
| | group_5842 | *PA3190* | *gltB* | | Q9HZ48 | Putative sugar-binding periplasmic protein | Carbon metabolism (pentose phosphate) | -5,31 | 1,43E-59 | -4,96 | 2,08E-23 | Activation | 7 |
| ▶ | edd | *PA3194* | *edd* | | P31961 | Phosphogluconate dehydratase | Carbon metabolism (pentose phosphate) | -2,16 | 1,26E-04 | -2,55 | 2,31E-10 | Activation | 7 |
| | epd_1 | *PA3195* | *gapA* | | P27726 | D-erythrose-4-phosphate dehydrogenase | Carbon metabolism (pentose phosphate) | -2,27 | 1,15E-05 | -2,42 | 1,87E-07 | Activation | 7 |
| | group_5870 | *PA3233* | *PA3233* | | Q9HZ07 | Hypothetical protein | Unknown | 2,04 | 1,84E-08 | 2,35 | 1,90E-05 | Repression | 7 |
| | actP_1 | *PA3234* | *yjcG* | s | Q9HZ06 | Cation/acetate symporter ActP | Unknown | 2,86 | 3,07E-07 | 2,89 | 5,65E-05 | Repression | 7 |
| | yjcH_1 | *PA3235* | *yjcH* | s | Q9HZ05 | Inner membrane protein YjcH | Unknown | 2,09 | 4,22E-04 | 2,31 | 6,97E-05 | Repression | 7 |
| | nosL | *PA3396* | *nosL* | | Q9HYK8 | Copper-binding lipoprotein NosL | Nitrogen metabolism | -2,63 | 2,89E-04 | -3,00 | 7,77E-03 | Repression | 9 |
| | mmsB | *PA3569* | *mmsB* | t | P28811 | 3-hydroxyisobutyrate dehydrogenase | Aminoacids (ILV) catabolism | 2,30 | 2,60E-10 | 3,27 | 1,43E-09 | | |
| ▶ | mmsA_1 | *PA3570* | *mmsA* | t | P28810 | Methylmalonate-semialdehyde dehydrogenase | Aminoacids (ILV) catabolism, propanoate and carbon metabolism | 2,32 | 8,63E-09 | 3,32 | 3,75E-08 | | |
| | iscR | *PA3815* | *iscR* | | Q9HXI7 | HTH-type transcriptional regulator IscR | isc operon regulation | -2,41 | 2,64E-06 | -2,10 | 3,56E-03 | | 13 |
| | dauA | *PA3863* | *dauA* | | Q9HXE3 | FAD-dependent catabolic D-arginine dehydrogenase DauA | Aminoacids (arginine, ornithine) catabolism | 2,12 | 6,53E-05 | 2,10 | 3,23E-03 | | |
| | argT_1 | *PA3865* | *PA3865* | | Q9HXE1 | Lysine/arginine/ornithine-binding periplasmic protein | ABC transporter (arginine, ornithine) | 2,47 | 2,45E-05 | 2,54 | 2,08E-04 | | |

| | | | | | | | | | | | | | |
|---|---|---|---|---|---|---|---|---|---|---|---|---|---|
| | group_3022 | ***PA3922*** | ***PA3922*** | u | Q9HX91 | Hypothetical protein | Unknown | **2,18** | 3,16E-07 | **2,13** | 8,05E-11 | | |
| | group_4203 | ***PA3923*** | ***PA3923*** | u | Q9HX90 | Hypothetical protein | Unknown | **2,39** | 2,29E-07 | **2,38** | 6,61E-11 | Activation | 9 |
| | group_2095 | ***PA4022*** | ***hdhA*** | | Q9HX05 | Hypothetical protein | Pyruvate and carbon metabolism, hydrazone utilization | **2,35** | 8,00E-12 | **2,50** | 4,71E-07 | | 27 |
| | yhdG_3 | ***PA4023*** | ***eutP*** | | Q9HX04 | Putative amino acid permease YhdG | Unknown | **2,79** | 7,48E-07 | **3,33** | 4,81E-08 | | |
| ► | acoR_1 | ***PA4147*** | ***acoR*** | | Q9HWN4 | Acetoin catabolism regulatory protein | Unknown | **2,02** | 1,07E-05 | **2,26** | 1,51E-05 | | |
| ► | fabG_10 | ***PA4148*** | ***PA4148*** | v | Q9HWN3 | 3-oxoacyl-[acyl-carrier-protein] reductase FabG | Butanoate metabolism | **2,52** | 7,65E-03 | **4,20** | 2,95E-08 | | |
| | acoA | ***PA4150*** | ***acoA*** | v | Q9HWN1 | Acetoin:2%2C6-dichlorophenolindophenol oxidoreductase subunit a | Unknown | **2,42** | 6,84E-03 | **3,89** | 5,15E-16 | | |
| ► | acoB | ***PA4151*** | ***acoB*** | v | Q9HWN0 | Acetoin:2%2C6-dichlorophenolindophenol oxidoreductase subunit b | Unknown | **2,95** | 8,68E-07 | **3,51** | 3,12E-15 | | |
| | acoC | ***PA4152*** | ***acoC*** | v | Q9HWM9 | Dihydrolipoyllysine-residue acetyltransferase component of acetoin cleaving system | Carbon and pyruvate metabolism | **3,00** | 4,18E-04 | **4,89** | 3,22E-19 | | |
| ► | ydjJ_2 | ***PA4153*** | ***PA4153*** | v | Q9HWM8 | 2,3-butanediol dehydrogenase | Butanoate metabolism | **2,33** | 9,28E-03 | **3,94** | 1,07E-13 | | |
| | pchA_2 | ***PA4231*** | ***pchA*** | | Q51508 | Salicylate biosynthesis isochorismate synthase | Ubiquinone and non-ribosomal siderophore peptides synthesis | **-2,72** | 3,55E-05 | **-2,18** | 5,58E-03 | | |
| | cckA | ***PA4293*** | ***pprA*** | w | Q9HWA7 | Sensor kinase CckA | Membrane permeability | **-2,54** | 2,98E-06 | **-3,28** | 1,55E-08 | Activation | 9,28 |
| | group_719 | ***PA4294*** | ***PA4294*** | w | Q9HWA6 | Hypothetical protein | Unknown | **-3,35** | 9,82E-14 | **-5,10** | 7,29E-14 | Activation | 9 |
| | group_2484 | ***PA4300*** | ***tadC*** | x | Q9HWA0 | TadC | Flp pilus assembly | **-2,45** | 8,38E-06 | **-2,73** | 4,22E-05 | Activation | 7,9 |
| | group_6095 | ***PA4301*** | ***tadB*** | x | Q9HW99 | TadB | Flp pilus assembly | **-2,47** | 1,84E-06 | **-2,73** | 1,06E-04 | | 31 |
| ► | group_4498 | ***PA4302*** | ***tadA*** | x | Q9HW98 | TadA ATPase | Flp pilus assembly | **-2,07** | 3,10E-05 | **-2,99** | 4,73E-06 | Activation | 7,9,29 |
| | outD | ***PA4304*** | ***rcpA*** | x | Q9HW96 | RcpA | Flp pilus assembly | **-2,48** | 2,53E-07 | **-2,92** | 1,79E-08 | Activation | 7,9,29 |
| ► | group_95 | ***PA4306*** | ***flp*** | | Q9HW94 | Type IVb pilin, Flp | Flp pilus assembly | **-2,94** | 1,73E-05 | **-2,67** | 1,65E-03 | Activation | 7,9,29 |
| | group_1582 | ***PA4638*** | ***PA4638*** | | Q9HVF3 | Hypothetical protein | Unknown | **-2,33** | 3,36E-08 | **-2,42** | 1,55E-04 | | |
| | yabJ_1 | ***PA5083*** | ***dguB*** | y | Q9HUA0 | 2-iminobutanoate/2-iminopropanoate deaminase | Aminoacids (glutamine) metabolism | **2,70** | 1,04E-04 | **3,77** | 1,29E-09 | | 30 |
| ► | dadA1_1 | ***PA5084*** | ***dguA*** | y | Q9HU99 | D-amino acid dehydrogenase 1 | Aminoacids (glutamine, phenylalanine) metabolism | **2,70** | 7,78E-05 | **4,01** | 2,78E-08 | | 30 |
| | group_5220 | ***PA5096*** | ***PA5096*** | | Q9HU87 | Glycine betaine-binding periplasmic protein OusX | ABC transporter (glycine, betaine, proline) | **2,51** | 2,24E-10 | **2,54** | 7,31E-06 | | |
| | proY_1 | ***PA5097*** | ***hutT*** | z | Q9HU86 | Proline-specific permease ProY | Aminoacids (histidine) catabolism | **2,46** | 1,71E-09 | **2,37** | 3,88E-07 | | 31 |
| | hutH_1 | ***PA5098*** | ***hutH*** | z | Q9HU85 | Histidine ammonia-lyase | Aminoacids (histidine) catabolism | **3,16** | 3,35E-16 | **2,95** | 2,39E-12 | | 31 |
| ► | pucI_1 | ***PA5099*** | ***PA5099*** | z | Q9HU84 | Putative allantoin permease | Aminoacids (histidine) catabolism | **2,84** | 6,41E-08 | **2,07** | 1,95E-05 | | 31 |
| | hutU | ***PA5100*** | ***hutU*** | | Q9HU83 | Urocanate hydratase | Aminoacids (histidine) catabolism | **2,65** | 4,00E-06 | **2,19** | 2,65E-08 | | 31 |
| | artJ | ***PA5153*** | ***PA5153*** | | Q9HU31 | ABC transporter arginine-binding protein 1 | ABC transporter (arginine) | **2,06** | 3,96E-05 | **2,23** | 8,34E-13 | | |
| | NA | ***PA5160.1*** | ***PA5160.1*** | | NA | tRNA-Thr(tgt) | Aminoacyl-tRNA biosynthesis | **3,26** | 5,20E-08 | **6,82** | 3,05E-20 | | |
| | group_4368 | ***PA5383*** | ***yeiH*** | | Q9HTI1 | Hypothetical protein | Unknown | **-2,93** | 3,32E-06 | **-3,25** | 1,16E-03 | | |
| | group_3039 | ***PA5460*** | ***PA5460*** | | Q9HTB0 | Hypothetical protein | Unknown | **-2,24** | 7,66E-09 | **-2,23** | 4,96E-03 | | |
| | group_5371 | ***PA5469*** | ***PA5469*** | | Q9HTA1 | Hypothetical protein | Unknown | **2,26** | 1,56E-03 | **3,02** | 4,99E-04 | | |

**Table S5: List of *P. aeruginosa* genes differentially expressed in presence of *S. aureus* in the context of coexistence.** PA2600 coexistence strain was cultivated in the absence or presence of SA2599 or SA2597. RNAs were extracted after 4 hours of culture and a RNAseq analysis was performed as described in material and methods. A gene was considered as differentially expressed when the Fold Change (FC) was > $|2\log_2|$ with an adjusted *P*-value<0.05 in presence of both SA strains. Genes from the same operon are annotated with an identical letter. Grey cells indicate genes that were also dysregulated in competition couples (Table S4). Symbol ► indicates genes tested in RT-qPCR (Fig. 1B). Functional classification was performed thanks to KEGG database and literature.

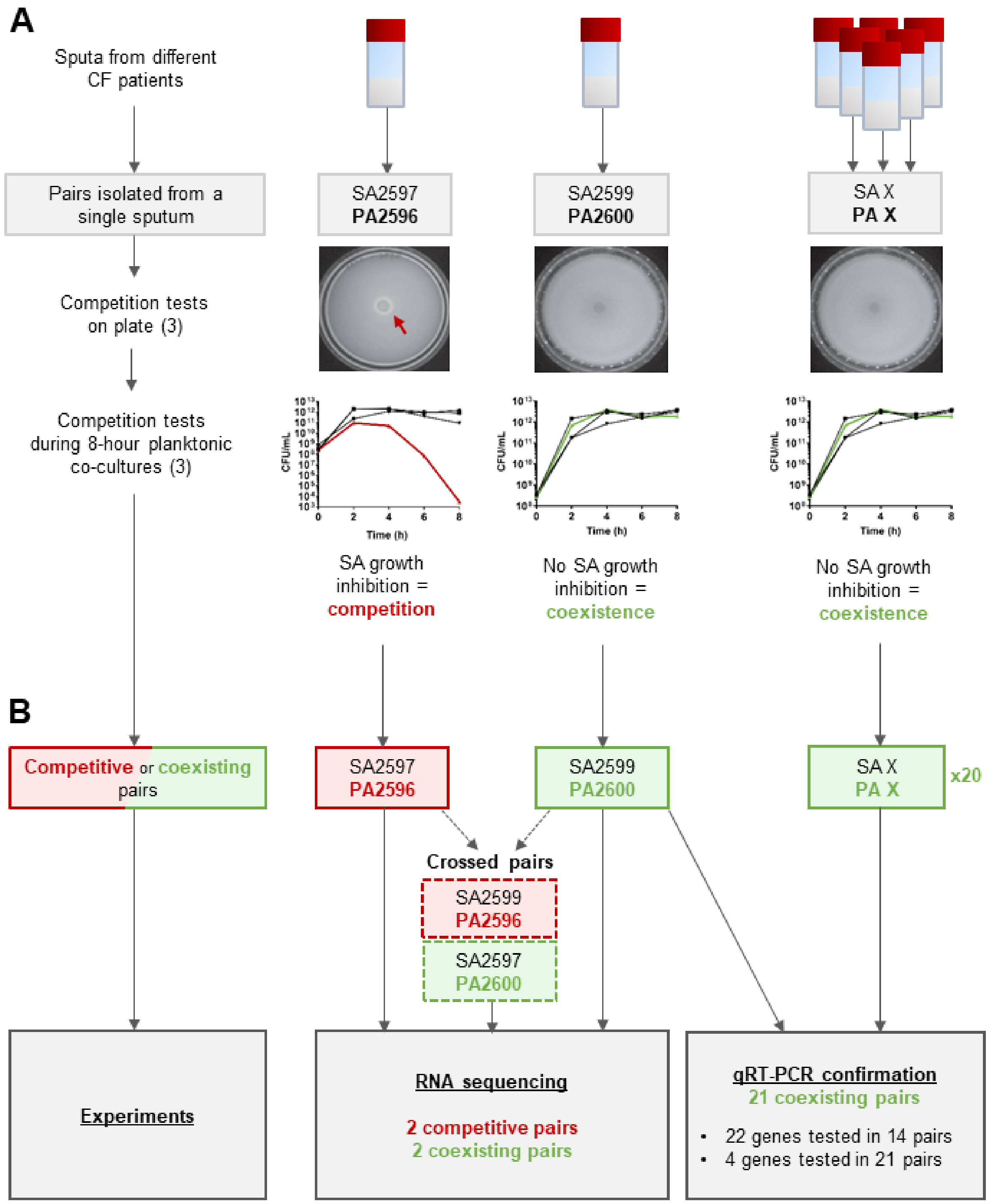


**Figure S1: Schematic representation of the employed methodology.**

**A. Determination of interaction state within *S. aureus*-*P. aeruginosa* co-isolated pairs.** Pairs of strains are co-isolated from a single sputum sample. Interaction state is tested during plate and liquid tests, as described in materials and methods and by Briaud *et al*. (3). Results of competition tests (pictures and kinetics) were obtained for a previous study (3).

**B. Strain pairs used in transcriptomic analyses.** Pairs SA2597/PA2596 and SA2599/PA2600 were isolated from two different patients. Interaction state of crossed pairs was determined as above and confirmed that it is solely led by *P. aeruginosa* (3). The 21 strain pairs used for qRT-PCR confirmation were both isolated from different patients, except in one case (Table S1).

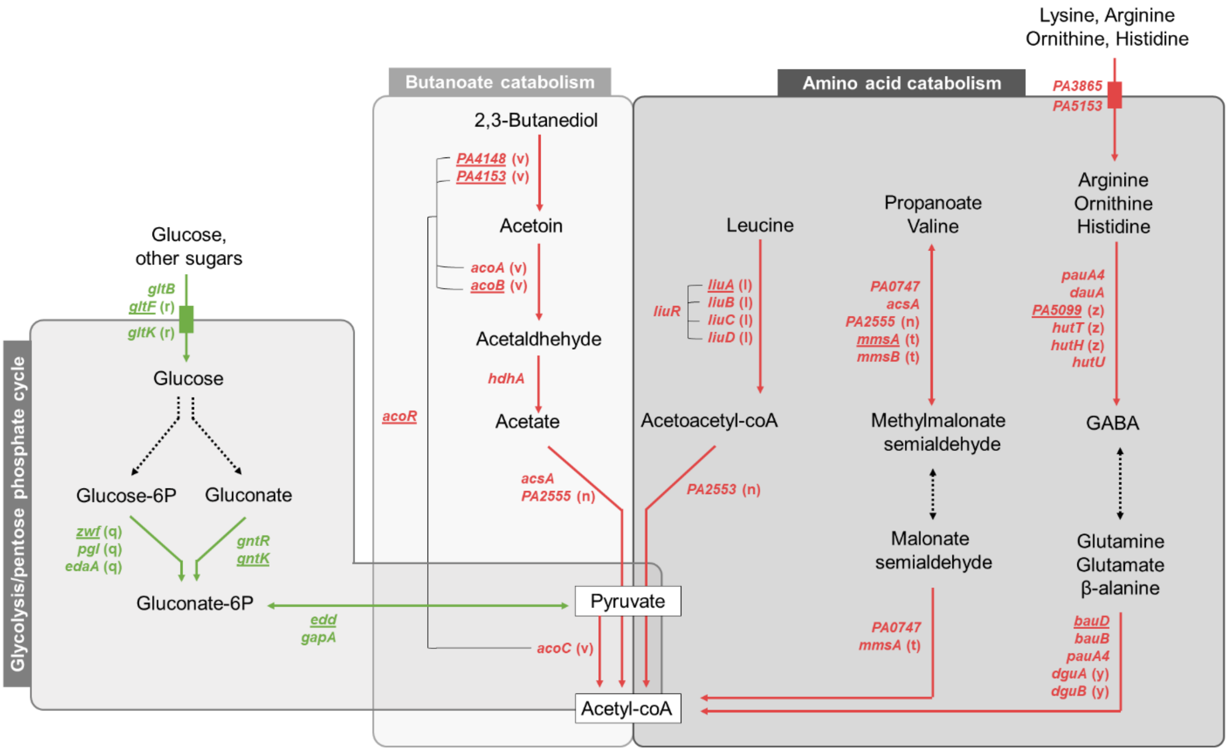


**Figure S2: *P. aeruginosa* metabolic pathways and associated genes up-regulated (red) or down-regulated (green) in coexistence with *S. aureus*.** PA2600 coexistence strain was cultivated in absence or presence of SA2599 or SA2597. RNAs were extracted after 4 hours of culture and RNAseq analysis was performed. A gene was considered as differentially expressed when the Fold Change (FC) was > $|2\log_2|$ with an adjusted *P*-value<0.05. Genes from the same operon are annotated with an identical letter. Genes tested in RT-qPCR and confirmed for PA2600 are underlined. Functional classification and pathway constructions were performed thanks to KEGG database and literature.

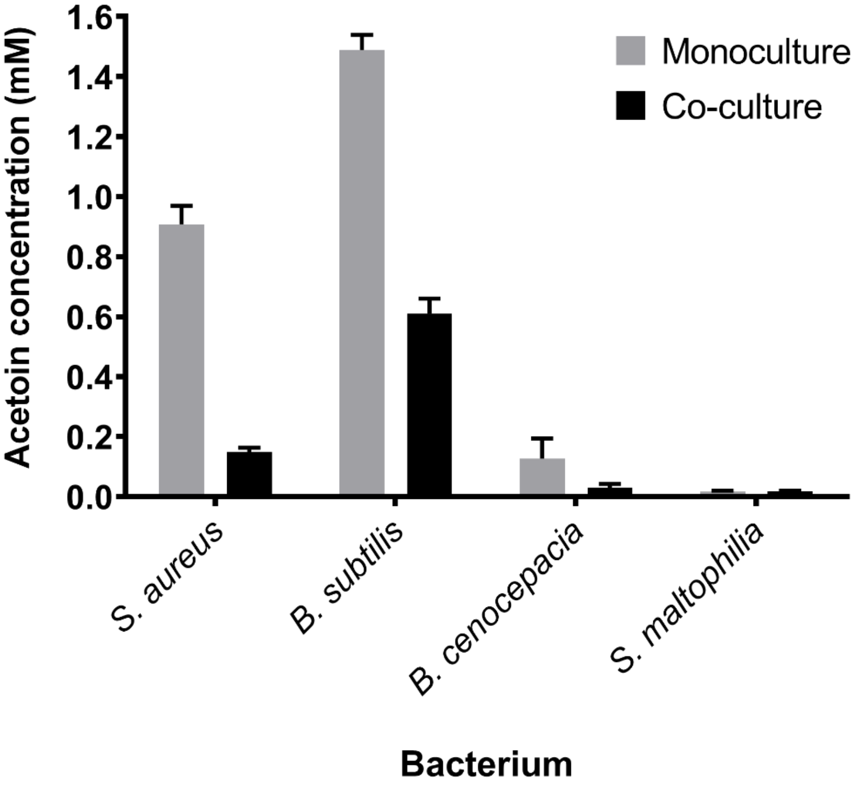


**Figure S3: Acetoin concentration in supernatant of *S. aureus* SA2599, *B. subtilis*, *B. cenocepacia* and *S. maltophilia* monocultures (grey bars) or co-cultures with *P. aeruginosa* PA2600 (black bars).** Acetoin was quantified from supernatant after 4h of culture. Bars represent the mean acetoin concentration + SEM from three independent experiments.

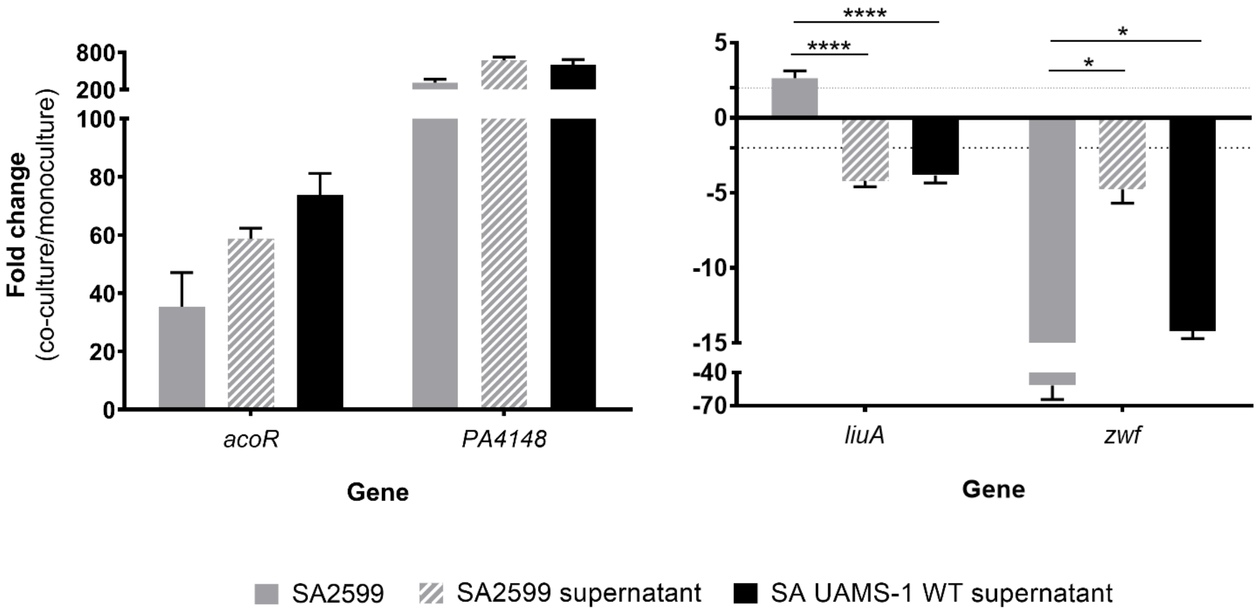


**Figure S4: Fold change of *P. aeruginosa acoR*, *PA4148*, *liuA* and *zwf* gene expression induced by culture with *S. aureus* (grey bars) or its supernatant (hatched and black bars).** *P. aeruginosa* PA2600 strain was cultivated in the absence or presence of *S. aureus* SA2599 or filtered supernatant of *S. aureus* SA2599 and UAMS-1 WT. RNAs were extracted after 4 hours of culture and gene expression was assayed by RT-qPCR. Bars represent the mean fold change + SEM from three independent experiments. Dot lines indicate a fold change = |2|. *$P_{adj}$<0.05, ****$P_{adj}$<0.0001 ANOVA with Tukey's correction.

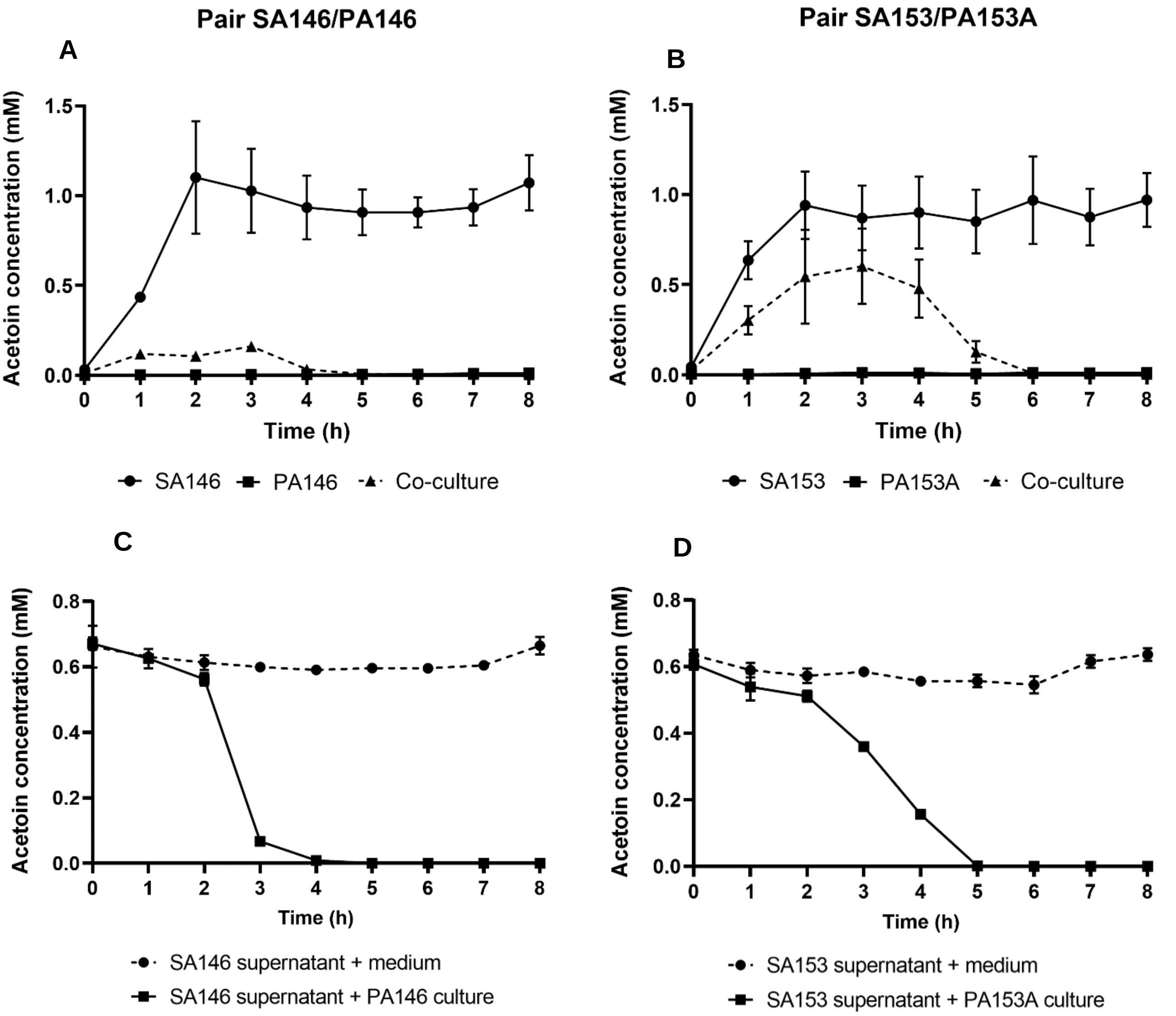


**Figure S5: Monitoring of acetoin concentration in *S. aureus* and *P. aeruginosa* monocultures or co-culture (A,B) or in *S. aureus* supernatant inoculated with *P. aeruginosa* (B,C), for the pairs SA146/PA146 (A,C) and SA153/PA153A (B,D).**

**A, B.** *S. aureus* and *P. aeruginosa* were cultivated in monoculture or co-culture. Acetoin was quantified from supernatant each hour. Points represent the mean acetoin concentration ± SEM from two independent experiments per pair.

**C, D.** A 4-hour filtered supernatant of *S. aureus* was inoculated with *P. aeruginosa* culture or sterile medium for controls. Acetoin was quantified from supernatant each hour. Points represent the mean acetoin ± SEM from three independent experiments per pair.

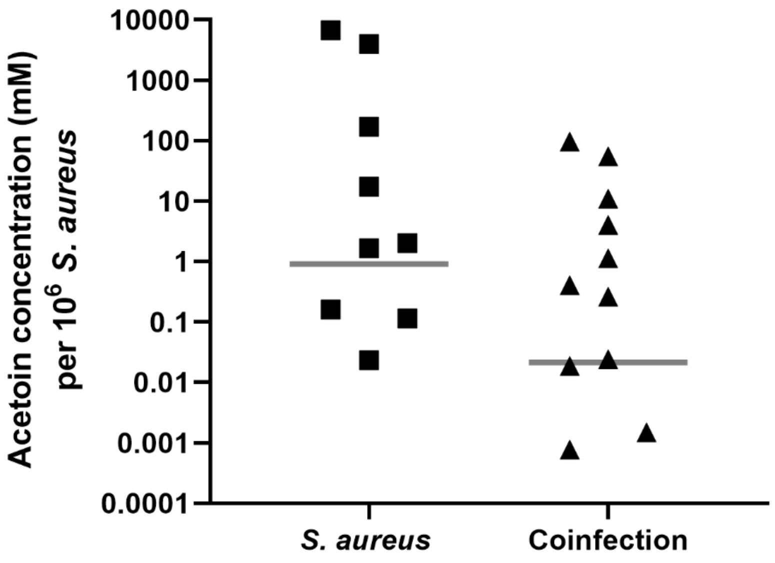


**Figure S6: Acetoin concentration in CF sputa from patients.** Sputa from *S. aureus* mono-infected patients (n=9) or *S. aureus* and *P. aeruginosa* co-infected patients (n=11) were gathered and acetoin concentration was quantified. Bars represent the median acetoin concentration normalized on *S. aureus* concentration in each sputum.

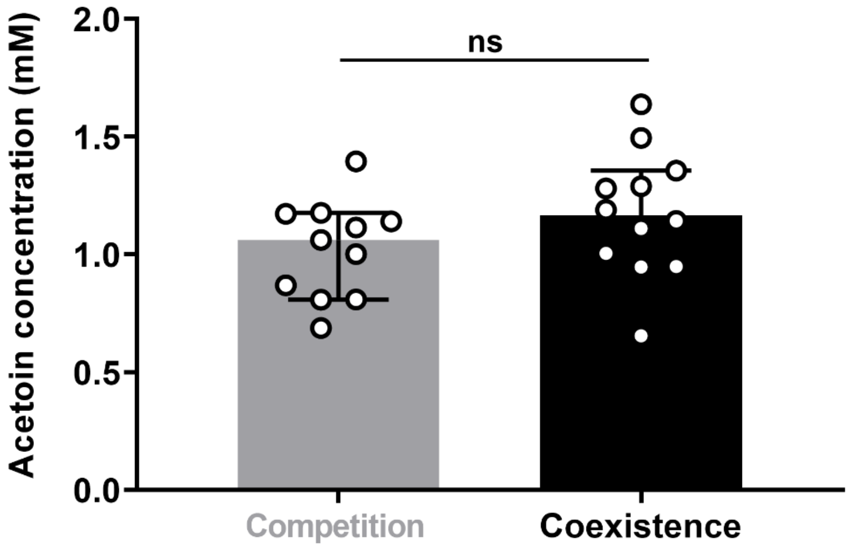


**Figure S7: Acetoin concentration in cultures of *S. aureus* strains from competition and coexistence couples.** Each *S. aureus* strain from competition (n=11) and coexistence (n=12) couples was cultivated for 6 hours in BHI and acetoin was dosed from supernatant. Bars represent the median acetoin concentration ± 95% CI. ns *P*>0.05 Mann-Whitney test.

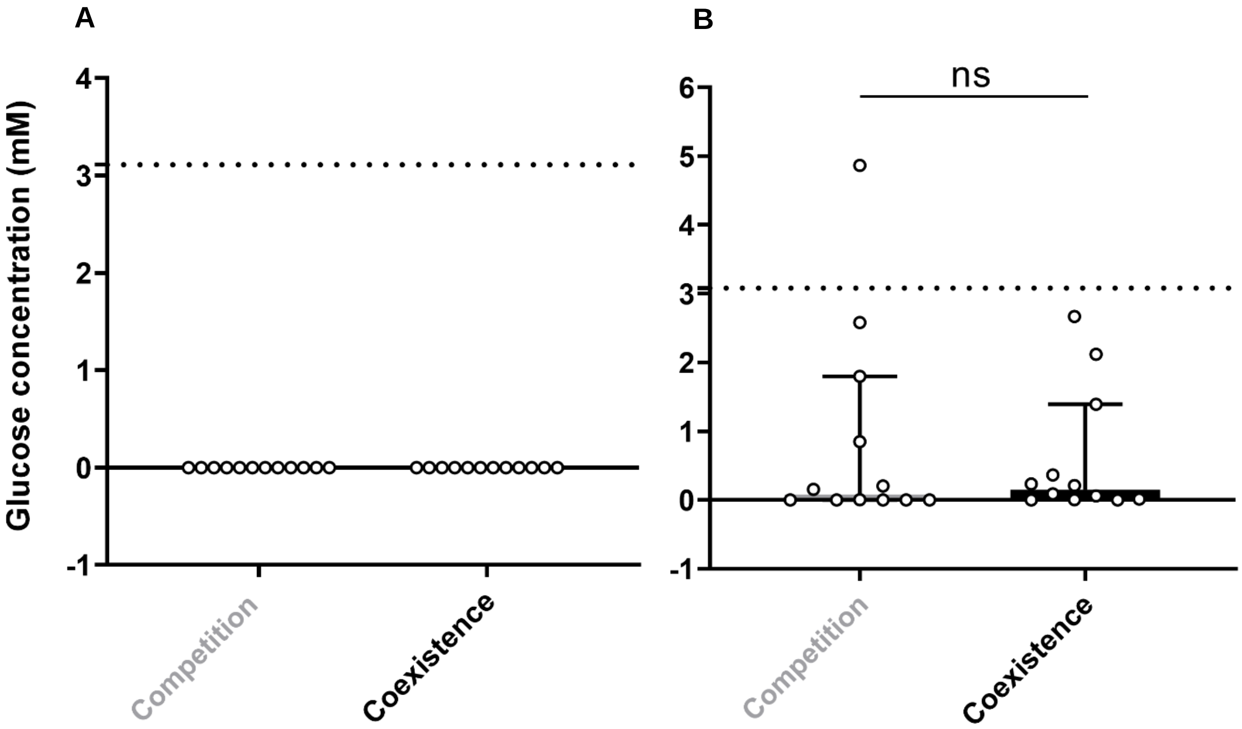


**Figure S8: Glucose concentrations in cultures of *S. aureus* (A) and *P. aeruginosa* (B) strains from competition and coexistence pairs.**

**A.** Each *S. aureus* strain from competition (n=12) and coexistence (n=12) couples was cultivated in *P. aeruginosa* PA2600 filtered supernatant for 6 hours and glucose was quantified from supernatant. No glucose was detected. Dotted line indicates the initial glucose concentration in *P. aeruginosa* supernatant.

**B.** Each *P. aeruginosa* strain from competition (n=12) and coexistence (n=12) couples was cultivated in *S. aureus* SA2599 filtered supernatant for 4 hours and glucose was quantified from supernatant. Bars represent the median glucose concentration ± 95% CI. Dotted line indicates the initial glucose concentration in *S. aureus* supernatant. ns *P*>0.05 Mann-Whitney test.

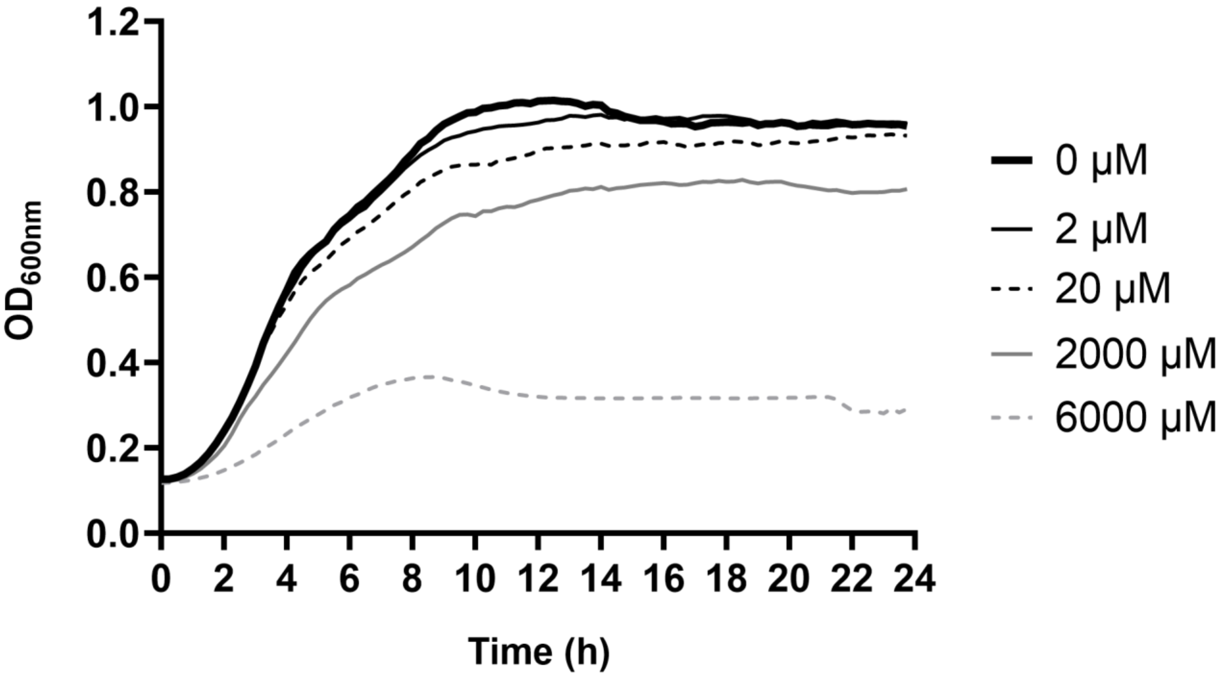


**Figure S9: Growth kinetic of *S. aureus* cultivated in absence or presence of acetoin.** SA2599 was cultivated during 24h in the absence of acetoin or in the presence of acetoin in different proportions ranging from 0.2µM to 6000µM per $10^6$ *S. aureus*. Lines represent the mean optical density of three technical replicates.

**Supplementary data references**

(1–5)(6–10)(11–15)(16–20)(21–25)(26–31)


1. Carriel D, Simon Garcia P, Castelli F, Lamourette P, Fenaille F, Brochier-Armanet C, Elsen S, Gutsche I. 2018. A Novel Subfamily of Bacterial AAT-Fold Basic Amino Acid Decarboxylases and Functional Characterization of Its First Representative: Pseudomonas aeruginosa LdcA. Genome Biol Evol 10:3058–3075.

2. Rietsch A, Vallet-Gely I, Dove SL, Mekalanos JJ. 2005. ExsE, a secreted regulator of type III secretion genes in Pseudomonas aeruginosa. Proc Natl Acad Sci U S A 102:8006–8011.

3. Briaud P, Camus L, Bastien S, Doléans-Jordheim A, Vandenesch F, Moreau K. 2019. Coexistence with Pseudomonas aeruginosa alters Staphylococcus aureus transcriptome, antibiotic resistance and internalization into epithelial cells. Sci Rep 9.
